# Artificial light at night consistently impacts avian physiology and behaviour: a meta-analysis

**DOI:** 10.1101/2025.10.17.683022

**Authors:** Diaz-Palma Sayuri, Capilla-Lasheras Pablo, Dominoni Davide, Cichoń Mariusz, Sudyka Joanna

**Affiliations:** Institute of Environmental Sciences, Faculty of Biology, Jagiellonian University, Poland; Doctoral School of Exact and Natural Sciences, Jagiellonian University, Kraków, Poland; Bird migration unit, Swiss Ornithological Institute, Sempach, Switzerland; Doñana Biological Station, Spanish National Research Council (EBD-CSIC), Sevilla, Spain; School of Biodiversity, One Health & Veterinary Medicine, University of Glasgow, UK

**Author notes:** **Email addresses:**,. **Corresponding author:** Sayuri Diaz-Palma, second corresponding author: Joanna Sudyka.

**Keywords:** artificial light at night, synthesis, avian performance, physiology, behaviour, life history traits, fitness, meta-analysis

## Abstract

Artificial light at night (ALAN) is a major anthropogenic pressure across the tree of life, driving population declines in insects and reptiles. Avian responses to ALAN defy simple patterns, varying in direction and magnitude, and the consequences of these responses remain unresolved. We conducted a meta-analysis to test how ALAN alters physiological, behavioural and life history traits underpinning avian performance. We analysed 623 effect sizes from 36 studies in 30 species. We found consistent physiological and behavioural shifts under ALAN, while life-history traits were unaffected. ALAN disrupted molecular and endocrine processes, leading to reduced sleep, higher metabolic rate, and accelerated reproductive maturation. Behaviourally, daily activity was extended, with earlier onset, later offset, and increased nocturnal activity and foraging effort. ALAN effects were stronger in migratory species and more pronounced in adults and females than nestlings and males. Higher light intensities amplified activity shifts, ageing, and sleep disruption, whereas exposure below 2 lux had minimal impact. Birds appear to buffer ALAN effects through physiological and behavioural adjustments, minimising impacts on life-history traits, a paradox likely explained by phenotypic plasticity or evolutionary adaptation. We identified light intensity threshold and group-specific vulnerabilities, emphasizing the need for targeted conservation in our increasingly illuminated world.

## INTRODUCTION

Artificial light at night (hereafter ALAN) has drastically transformed the predictable natural light-dark rhythms on Earth. In just the past decade, ALAN has been increasing by approximately 2.2% annually and has been emerging as a major source of light pollution and a pronounced anthropogenic disturbance (Falchi *et al*. 2016; Gaston *et al*. 2014). Negative effects of ALAN have been documented across a wide range of taxa, including plants (Bennie *et al*. 2016; Matzke 1936), microorganisms (Cesarz *et al*. 2023; Pu *et al*. 2019), marine invertebrates (Duarte *et al*. 2019; Underwood 2024), fish (Pulgar *et al*. 2023; Schligler *et al*. 2021), amphibians (Touzot *et al*. 2023; Wise 2007), reptiles (Kolbe *et al*. 2021; Taylor *et al*. 2022), birds (Adams *et al*. 2019; Xue *et al*. 2020), and mammals (Ditmer *et al*. 2021; Shier *et al*. 2020). ALAN affects organisms through multiple pathways: disrupting natural light-dark cycles that entrain circadian rhythms governing the physiology and behaviour (Bumgarner & Nelson 2021; Cassone & Kumar 2022), and causing direct mortality through attraction, collision and increased predation. While attraction-related deaths clearly impact population persistence (Desouhant *et al*. 2019; Grunsven *et al*. 2020; Owens *et al*. 2020), whether circadian disruption independently drives biodiversity loss remains uncertain (Jägerbrand & Spoelstra 2023).In animals, attraction to ALAN is a well-known direct response leading to reduced fitness and population declines, but the available literature remains restricted to a limited number of taxa, including insects, sea turtles, and birds (Leader *et al*. 2024; Rodriguez *et al*. 2017; Salmon 2003; Van Doren *et al*. 2021). A recent meta-analysis found no global population or community effects of ALAN across taxa (Sanders *et al*. 2020) but evidence for population declines is clear in specific groups. Insects suffer from both fatal light attraction and behavioural and physiological disruption (Boyes *et al*. 2021) and sea turtle populations decline primarily through light-induced hatchling disorientation rather than circadian disruption (Dimitriadis *et al*. 2018). Birds exemplify a more complex pattern. Despite showing pronounced ALAN-induced responses in sleep, behaviour, cognition and other mechanisms underlying fitness (Aulsebrook *et al*. 2021; Dominoni 2015), population-level impacts vary-some species decline while others persist or even thrive in illuminated environments. This is particularly striking given that birds, unlike most vertebrates, are often abundant in urban environments where nocturnal light is prevalent (Aulsebrook *et al*. 2021; Dominoni 2015), making them exemplary models for understanding variation in ALAN responses across biological levels (Aulsebrook *et al*. 2021; Bumgarner & Nelson 2021).

A growing body of evidence suggests that light pollution may significantly alter specific aspects of avian performance (Dominoni 2015; Sepp *et al*. 2018). Avian performance reflects an interplay of physiological, behavioural, and life history-traits that determine a bird’s capacity to maintain homeostasis with its environment (Kempenaers 2022; Sibly *et al*. 2012; Wingfield *et al*. 2017). Within these categories, specific functional traits, which are defined here as measurable characteristics of biological relevance, may respond directly to ALAN (Nock *et al*. 2016). The primary mechanism underlying these responses is disruption of the circadian clock, which orchestrates biological functions according to natural light-dark cycles (Cassone 2014; Rock *et al*. 2022). This disruption interfaces with immunoregulation, stress receptors and metabolic pathways (Gotlieb *et al*. 2018; Jerigova *et al*. 2022; Markowska *et al*. 2017) and affects melatonin secretion, which modulates sleep, metabolism, and activity in diurnal birds (Cassone 2014; Gwinner *et al*. 1997; Wang *et al*. 2012). Moreover, ALAN responses vary across life stages, sexes, and ecological strategies. For instance, nestlings faced high energy demands and greater physiological sensitivity than adults, enforced by the fact that unlike many young mammals, must develop rapidly to fledge (Assadi & Fraser 2021; Grunst *et al*. 2020; Oswald *et al*. 2018; Vézina & Salvante 2010). At adulthood, environmental stressors may differentially impact reproductive energy allocation between sexes, with males investing in foraging and females facing greater endocrinological challenges (Frank 1990; Sibly *et al*. 2012; Tarwater & Arcese 2017). Migratory behaviour and habitat preferences (e.g., open vs dense habitats) have also been shown to influence avian vulnerability to anthropogenic impacts such as ALAN (Adams *et al*. 2019; Bhagarathi *et al*. 2024; Burt *et al*. 2023). Consequently, reported effects vary widely across species and contexts.

ALAN affected physiological traits such as immune function, with dark-eyed juncos showing increased blood parasite loads (Becker *et al*. 2020) and caused sleep dysregulation via reduced melatonin levels in adult great tits, as well as earlier reproductive activation through accelerated testes growth (Dominoni *et al*. 2018; de Jong *et al*. 2016). Behaviourally, ALAN consistently extended daily activity and vocal periods across species (Champenois *et al*. 2025; Dominoni *et al*. 2014, 2022; Graham *et al*. 2019; Pease & Gilbert 2025), yet sleep patterns appeared context-dependent, occurring for example in breeding female great tits under specific light intensities (Raap *et al*. 2016b; Ulgezen *et al*. 2019). Notably, findings for life-history traits reveal marked inconsistencies. Most studies reported no effects on reproductive output, such as clutch or brood size (Ferraro *et al*. 2020; Grunst *et al*. 2019; de Jong *et al*. 2015; Kempenaers *et al*. 2010; Ouyang *et al*. 2015) and some documented reduced hatching success and delayed phenology (Assadi & Fraser 2021; Champenois *et al*. 2025; Injaian *et al*. 2021; Malek & Haim 2019). Similarly, body mass responses vary; several studies reported no changes (Ferraro *et al*. 2020; Injaian *et al*. 2021; Kumar *et al*. 2021; Moaraf *et al*. 2021), whereas others attributed nestling weight loss in response to ALAN to reduced parental care (Cianchetti-Benedetti *et al*. 2018; Reid *et al*. 2025) and increased energy expenditure (Ferraro *et al*. 2020; Raap *et al*. 2016a).

Overall, the impact of ALAN on birds has been extensively documented with reviews examining endocrine and behavioural mechanisms (Dominoni 2015; Grunst & Grunst 2023; Helm & Liedvogel 2024) or ecological outcomes (Adams *et al*. 2021; Jägerbrand & Spoelstra 2023). Sanders et al., 2020 provided a valuable synthesis of ALAN’s effects across taxa, including birds, identifying effects on physiology, activity patterns, and life-history traits. Although they also disentangled these effects within subcategories such as gene expression and hormone levels, these measures encompass a variety of functional traits. While ALAN clearly alters diverse functional traits, the direction and strength of effects on avian performance remain unclear, varying within and across species depending on life stage, sex, and environmental context. Understanding which functional traits underpinning avian performance are most sensitive to light pollution is essential for conserving biodiversity, guiding urban mitigation strategies and advancing ecological and evolutionary research. Towards this goal, we conducted a systematic review and a meta-analysis to quantitatively synthesize the effects of ALAN on avian performance. Based on previous studies, we selected specific moderators to investigate whether ALAN affects avian functional traits related to physiology, behaviour, and life history traits, and to evaluate how these effects vary within and between species. The three broad categories of avian performance were divided to 16 functional traits covering various measurements of intra-specific (e.g., life stages and sexes) and inter-specific traits (e.g., migratory status, habitat densities, study environment (lab vs. field) and population trends). Specifically, we tested whether ALAN was associated with disruption of circadian clock functioning by shifting the expression of core clock genes, particularly at night, with endocrine-molecular downstream effects on ageing, immunity, metabolic rate, neuronal cognition, sleep regulation and reproductive maturation. These physiological imbalances could affect aspects of behaviour, such as activity offset and onset, foraging effort, levels of daily and nocturnal activity, and ultimately life-history related traits (body mass and size alongside reproductive phenology and success). Additionally, we assessed ALAN’s effects under varying experimental regimes (light intensities, days since ALAN exposure, day vs night and tissue types) to identify the specific conditions under which ALAN exerts the strongest impact on avian performance.

## METHODS

### Literature review

We performed a systematic research of published literature in three different databases: Scopus, Web of Science (All databases) and PubMed on the 28^th^ of February 2023 through an institutional subscription via the Jagiellonian University, Krakow, Poland. We retrieved relevant literature on the effects of ALAN on avian performance using the following search string: (artificial light OR light* pollut* OR light* at night OR night time light* OR alan OR lux OR lx OR light* disturb* OR city light* OR urban* light*) AND (bird* OR avian OR feather*) AND (circadian clock* OR rhythm* OR period* OR physiolog* OR metabol* OR hormon* OR melatonin* OR gene* OR immun* OR perform* OR fitness* OR life-history* OR surviv* OR phenotyp* OR body size OR body condition OR activit* OR sleep* OR behav* OR song* OR sing* OR flight* to light OR reproduct* OR breed* OR mating* OR incubat* OR predat* OR develop*) AND (adult* OR young OR nestling* OR chick* OR hatch* OR egg* OR migra*) NOT (broiler*). The search yielded 1,480 records from which we included 27 studies more that were reported in a previous meta-analysis (Sanders et al., 2020) and fitted our inclusion criteria. In total 1,507 records were imported into Rayyan (Ouzzani *et al*. 2016). After removing duplicates (n=388), 1,128 unique studies remained and were selected for screening. The results of each search phase are reported in the Preferred Reporting Items for Systematic Reviews and Meta-Analysis (PRISMA) diagram (supplementary material, Fig. S1).

### Inclusion Criteria

The first screening consisted in reading the title and abstract to include studies using the following criteria: (1) studies reporting effects of ALAN on avian species in the field or laboratory; (2) including quantitative measurements of light (measured in lux); (3) reporting ALAN-DARK paired comparisons, where the DARK condition corresponds to a control group exposed to natural nocturnal light levels (≤ 0.2 lux), and the ALAN group is exposed to ALAN up to 100 lux. We considered 0.2 lux as the maximum natural night illumination, as lunar light intensity can reach up to 0.26 lux during the super moon occurring once per year (Kyba *et al*. 2017). As for ALAN light intensity levels, we aimed for the studies reporting realistic real-life values; street lamps produce on average 1.6 10 lux, with urban light pollution reaching maximal intensity of 100 lux on an illuminated stadium (Gaston *et al*. 2013; Rich & Longcore 2013; Sanders *et al*. 2020). We excluded literature reviews, conference reports and patents. We also removed studies concerning meat/egg production or any pedigree-bred birds to avoid including organisms artificially selected for specific traits, which could potentially influence their response to ALAN. Studies related to photoperiod, natural day light and studies comparing effects of light from urban areas were also omitted to avoid a possible influence of external disturbance and confounding factors such as noise or human activity (Wilson *et al*. 2021). Finally, we excluded data from within individual design studies, where the same group of birds was exposed to both ALAN and dark conditions at different time points. We took this decision to homogenise the experimental designs of the studies included this meta-analysis and to avoid the potential influence of prior exposure to ALAN in the response to natural dark conditions (and vice versa). The first screening was repeated in 10% of the studies by P. C-L. and J.S., resulting in 95% of inclusion consistency. A description of the number of studies excluded, and the exclusion reason is shown in Supplementary material (Table S7).

### Full screening and data extraction

The first screening yielded 171 studies that we read fully to extract the quantitative data needed to calculate effect sizes for ALAN–DARK paired comparisons. For each study, we collected the following moderators: publication year, life-stage (adults or nestlings), sex, study environment (captivity or wild), focal species and light intensity. We aimed to extract mean, standard deviation (SD), and sample size (n) from both groups. Our final dataset comprised 36 studies, from which we extracted 65 effect sizes from tables and text, and 192 effect sizes from raw data provided in the electronic supplementary materials. Since we focused on estimating the main effect of ALAN on avian performance traits, we excluded data from models reporting interactions (e.g., sex or age), unless the authors provided data to extract individual measures from them. We used ‘Web Plot Digitizer’ software (Tantry *et al*. 2021) to extract means and SD from figures when these values were not provided in the text or tables (n = 360 effect sizes). We calculated standard errors for 13 effect sizes from 2 different studies, which were reported as 95% confidence intervals (Cooper *et al*. 2009). We used the formula: SD = SE× √*n* to calculate SD from 285 standard errors (SE). In three studies, we calculated the mean and SD from eight quartiles and medians following (Hozo *et al*. 2005). From four studies, we extracted the mean and SD from 17 estimates from linear mixed models reporting differences between ALAN and DARK groups (the method and formula are reported in the supplementary material text). We confirmed that effect sizes derived from raw data did not differ significantly from those obtained in models by including type of effect size (raw vs model) as a moderator. As type of effect size did not account for a significant proportion of variance, the variable was excluded from the final models. When the data reported in a study were incomplete (e.g. the mean was given, but the SD, SE, or sample size for one of the groups was missing), we contacted the authors. Five of the fourteen authors we contacted, everyone who provided additional information or raw data, are mentioned in the acknowledgements.

### Mapping effect sizes onto avian performance functional traits

To provide a comprehensive framework for understanding avian responses to ALAN, we assigned the effect sizes to 16 functional traits underpinning physiological, behavioural and life history mechanisms of avian performance. Physiological traits included ageing, elements accelerating (telomere length) or slowing (antioxidants) physiological decline over time; circadian clock functioning, as the expression of core clock genes; immunity, defence responses against pathogens (e.g., anti-bacterial response); metabolic rate, factors contributing to energy production and expenditure; neuronal cognition, mechanisms underlying cognitive functions (e.g. neuronal density); reproductive maturation, processes contributing to reproductive development and activation (e.g. gonadal maturation); and sleep regulation, referring to biochemical modulators of sleep onset and duration (e.g. melatonin levels). Behavioural functional traits comprised activity offset, as the time of end of daily activity; activity onset, as the time of start of daily activity, foraging effort, the investment in food-searching and handling (e.g., preys per minute or food intake); level of activity (e.g., activity counts); and nocturnal activity (e.g., activity counts at night). Finally, life history traits encompassed body mass and size, as any phenotypical measure related to individual condition and fitness maintenance; reproductive phenology, as any seasonal timing of reproductive events (e.g., fledging date); and reproductive success, the outcome of breeding efforts (e.g., number of fledglings). We provide a detailed description of each functional trait in Table 1.

**Table 1.** The table shows 16 avian functional traits underpinning avian performance, sorted by meta-analytical categories: physiology (red), behaviour (blue) and life history (violet). Functional traits are presented with a ‘functional note’ describing their relevance in the context of avian performance. ‘Example proxies’ include the most representative variables from the published literature assigned to each trait. Variables expected to negatively impact the corresponding functional trait are marked with (-). The full list of effect size assessments is shown in Supplementary Table S1.

| Level | Avian functional trait | Functional note | Example proxy |
| --- | --- | --- | --- |
| Physiology | Ageing | Decline in physiological integrity over time affecting reproduction and survival | Relative telomere length, TAC- total antioxidant activity (-) |
|  | Circadian clock functioning | clock core genes regulating physiology and behaviour within 24h light-dark cycle | Nocturnal Bmal1, Per2 mRNA expression (-) |
|  | Immunity | Pathogen defence; vital for health and fitness | HP- haptoglobin, % bacterial E. colli killing |
|  | Metabolic rate | Biochemical processes for energy production and allocation | Daily energy expenditure (DEE), Body temperature |
|  | Neuronal cognition | Biochemical mechanisms underlying cognitive functions | Neuronal densities linked to neuronal plasticity |
|  | Reproductive maturation | Endogenous timing, activation and development of reproduction | Luteinizing hormone (gonadal maturation) |
|  | Sleep regulation | Biochemical modulators of sleep onset and duration, vital for behavioural functioning | Melatonin- induce sleep onset, achm3 - awake promoter gene (-) |
| Behaviour | Activity offset | Time of day when activity ends | Last activity of the day and parental visit |
|  | Activity onset | Time of day when activity begins | First nest entry or chorus time |
|  | Foraging effort | Food acquisition strategies | Food intake, and preys per minute |
|  | Level of activity | Behaviours duration linked to energetic expenditure | Activity duration and activity counts per hour |
|  | Nocturnal activity | Extent of behavioural activity during night-time hours | Sleep duration- less activity (-), Awakening time |
| Life history traits | Body mass | Indicator of energetic reserves supporting survival and reproductive investment | Body mass, weight and growth rate |
|  | Body size | Body dimensions shaping metabolism and reproductive capacity | Tarsus, wing, and rectrices lengths |
|  | Reproductive phenology | Seasonal reproductive timing to match with favourable conditions | Incubation, nesting, and fledging dates |
|  | Reproductive success | Outcome of breeding efforts directly related to fitness | Parental feeding rate, Nest predation (-) |

To minimize the risk of inconsistently labelling the effect sizes into functional traits, we based the assignment on the definitions provided in the original studies. Since some genes and physiological features may capture diverse biological functions, each effect size was assigned to a specific functional trait based on the main function targeted in the original study. To enable interpretation of the meta-analysis results, effect sizes expected to have a negative impact on the corresponding functional trait were multiplied by -1, unless the sign was already provided in the original study. We included effect sizes for reproductive output only from the first brood. Effect sizes reported across a 24-hour period were categorized as either ‘day’ (e.g., morning, noon) or ‘night’ (e.g., midnight) traits. We calculated the difference in mean and SD from eight studies (n= 67 effect sizes) reporting effect sizes before (baseline) and after ALAN exposure (details of the conversions are provided in the supplementary material). We included functional traits in the global meta-analytical model only if they were reported in at least three different studies. Circadian clock functioning was reported in only two studies and then excluded from the main model. A description of the assignment of effect sizes to functional traits and their directionality is provided in Supplementary material, Table S1.

### Phylogeny and inter-species variation

To account for the phylogenetic variation in our meta-analytic models, we computed a correlation matrix based on species relationships. This matrix was derived using branch lengths from Open Tree of Life phylogenies via the ‘rotl’ R package trees (https://opentreeoflife.github.io) (Michonneau *et al*. 2016). The phylogenetic tree used for the global analysis is shown in Fig. 1B. We also retrieved the migration status (migratory, partially migratory, or sedentary) and habitat density (dense, open, or semi-open) from the AVONET database (Tobias *et al*. 2022), and the population trend (decreasing, increasing, or stable) from the IUCN Red List of Threatened Species™ (https://www.iucnredlist.org) for each species. A summary of intra-specific traits is shown in supplementary material, Table S2.

**Figure 1.**
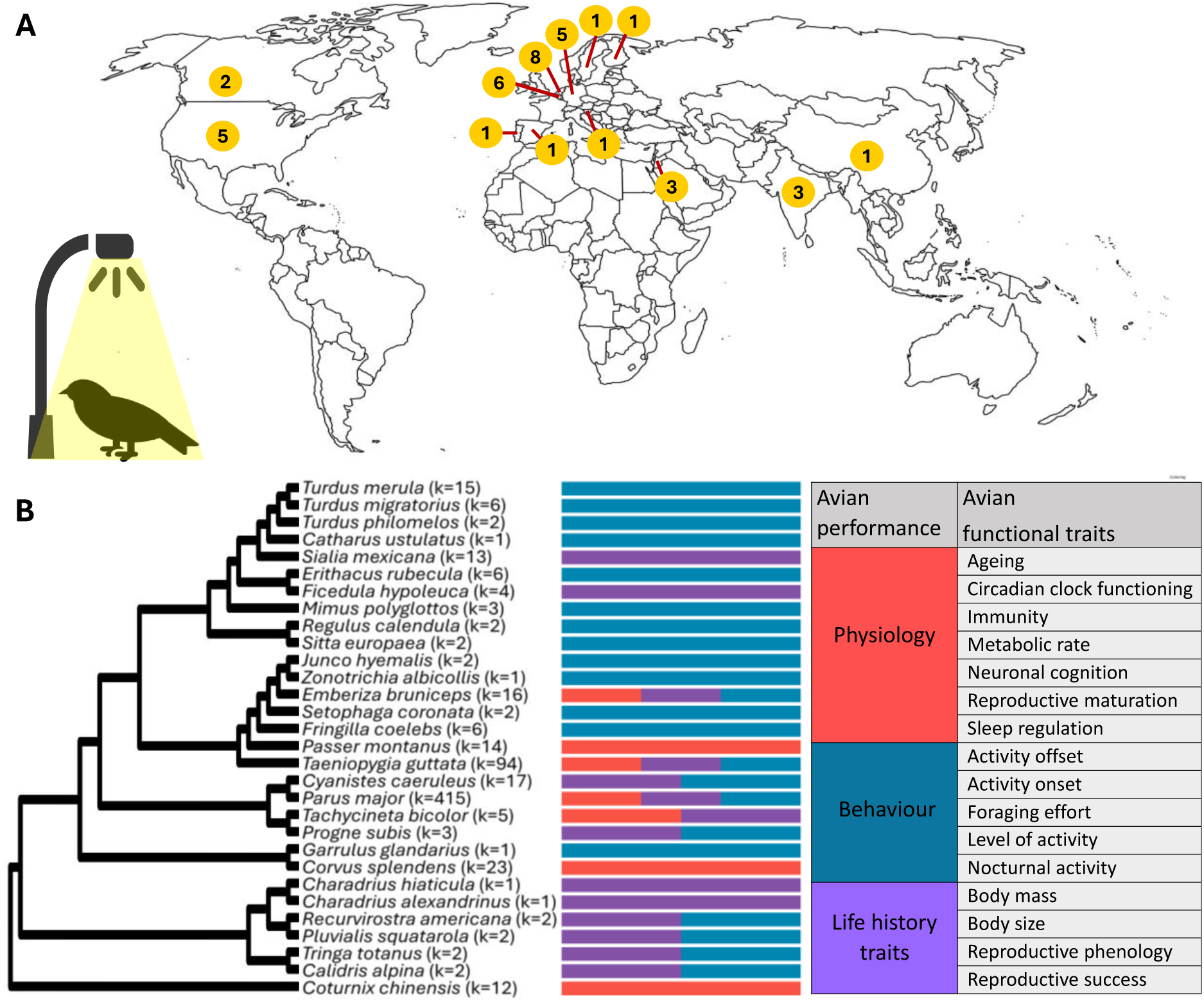
Geographic and phylogenetic distribution of the meta-analytical dataset including 675 effect sizes reporting effects of artificial light at night on avian performance. A) Map showing the distribution of the 36 studies included in the meta-analysis. Numbers in yellow indicate the number of studies per country. B) Phylogenetic tree of the 30 avian species included in the meta-analytical model. For each species, k represents the number of effect sizes and bars show the proportion of effect sizes assembled in the following categories of avian performance: physiology (red bars), behaviour (blue bars) and life history traits (purple bars). The table shows the 16 functional traits sorted by avian performance category included in the meta-analysis.

### Effect sizes

We calculated standardized mean differences between paired ALAN and DARK effect sizes. We used the function ‘escalc’ in ‘Metafor’ R package (v. 4.6 ; R Core Team, 2024) to calculate Hedges’s g and the sampling variance. Our final dataset included 675 ALAN-DARK paired comparisons from 30 species reported in 36 different studies published between 2006 and 2022.

### Data analysis

All the models of this meta-analysis were conducted in R (v.4.2.2; R Core Team, 2022), using the package ‘metafor’ v.4.6-0 (Viechtbauer 2010). First, we fitted a phylogenetic multi-level (intercept-only) meta-analysis with Hedges’ *g* as the response variable. This model allowed us to estimate heterogeneity in ALAN effects across the dataset, including within-study, between-study, and phylogenetic variance components, and to provide the overall mean effect size across traits (Table 2; Model 1). Then, we assessed the directional influence of ALAN on each functional trait by including the functional trait category as a moderator (Table 2; Model 2). Second, we evaluated the influence of intra-specific variation by comparing effects of ALAN on functional traits between life stages (adults vs. nestlings) and sexes (males vs. females). To enable direct comparisons, we ran separate models for adults (Table 2; Model 4) and nestlings (Table 2; Model 5) from subsets containing the same avian functional traits, and we followed the same approach for females (Table 2; Model 6) and males (Table 2; Model 7). Third, we tested the influence of inter-species variation by fitting a multi-level model with migration and habitat density as moderators (Table 2; Model 8). Since we observed moderate collinearity among moderators (GVIFs < 5) in this model, we used Type II tests, which evaluate each moderator’s significance while accounting for the others. Then, because species traits were not available for all the species, we tested the influence of study environment and population trend in separate models (Table 2; Models 9 and 10). All the models testing the effects of intra and inter-specific traits included avian functional traits as moderators.

**Table 2.**
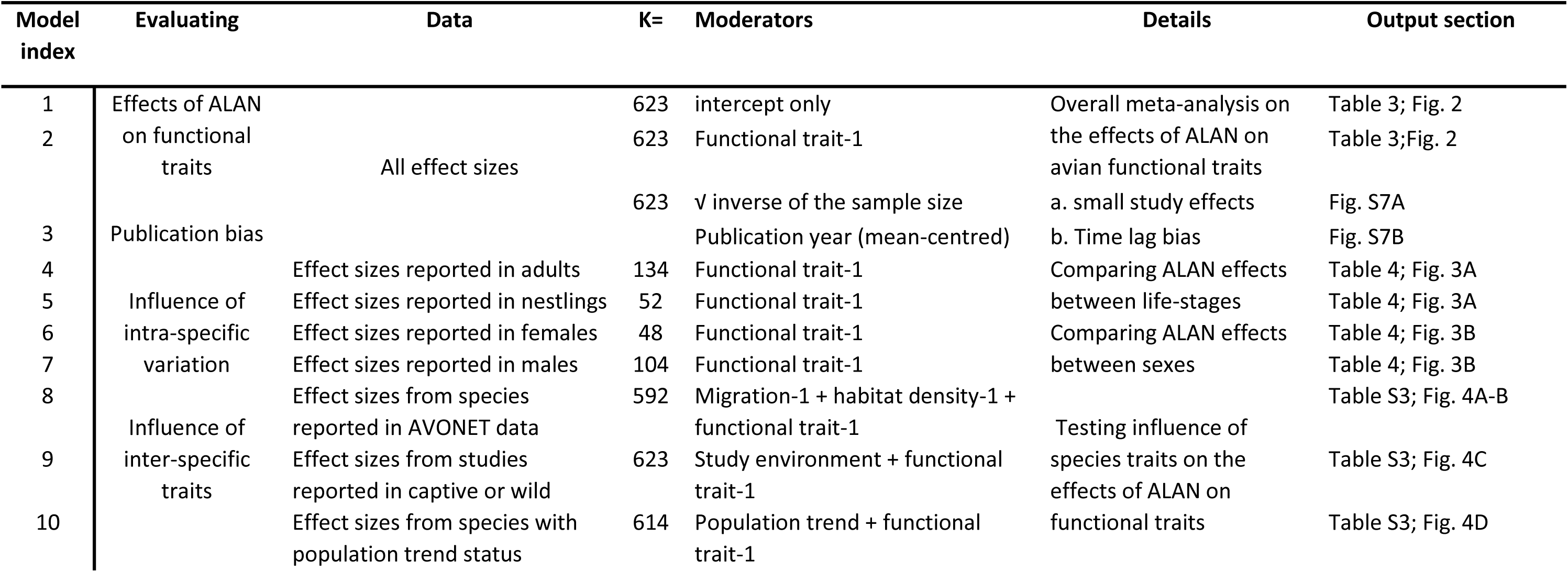
Description of the meta-analytical models evaluating the effects of light pollution on avian performance. ‘Model index’ contains a series of sequential numbers from 1 to 10 to facilitate the understanding of the methods and results. ‘Data’ refers to the effect size inclusion and k represents the number of effect sizes for each model. ‘Moderators’ show the categories included as moderators in each model, ‘Details’ provide a short description of the model and ‘Output section’ show where to find the model output.

We accounted for among and within study variation in every model by including study ID and individual observation ID as random effects. We included species ID and the phylogeny as a random effect when the model included two or more different species. We used the ‘i2_ml’ function from the OrchaRd package (v. 2.0) to calculate the percentage of total relative heterogeneity and the heterogeneity explained by each of the random factors (Nakagawa *et al*. 2023).

We conducted additional subset analyses. Since we consider the circadian clock fundamental to all physiological, behavioural, and life-history changes induced by ALAN, we tested the effects of ALAN on traits associated with circadian clock function by running an intercept-only model and then models including light intensity and time of day as moderators. Given that light intensity was reported for most effect sizes, we evaluated its influence across overall functional traits. Then, we tested the influence of varying experimental regimes included in our dataset: light intensities (e.g., 0.5, 1, 5 lux), days since ALAN exposure, time of day (day vs. night), and tissue types (e.g., liver, spleen, brain) by grouping repeated effect sizes per condition. Individual subset analyses were then conducted for each functional trait. A summary of the models included in the subset analysis is shown in supplementary material, Table S4; Models 11-15.

### Testing for publication bias

We examined the data for indicators of two types of publication bias, small study effects and time lag bias, following Nakagawa et al. (2023). We ran two additional multilevel meta-analytical models including as a single moderator either the square-root of the inverse of the effective sample size or the mean-centred year of study publication (Table 2; Model 3). We used the ‘r2_ml’ function from the orchaRd package (v. 2.0) to calculate the variation explained by these moderators (R^2^_marginal_) for each model (Nakagawa et al., 2022).

## RESULTS

### Effects of ALAN on avian functional traits

The intercept only model revealed a nonsignificant near-zero effect of ALAN on the overall effect sizes (mean estimate [95% CI] = 0.056 [-0.482, 0.593]; Fig. 2). Total heterogeneity was high (I^2^ = 89.9%), with most variance arising within studies (57.6%), followed by among studies (16.5%) and phylogenetic effects (14%), while among-species heterogeneity was low (1.8%). Model 2 (Table 3) including functional traits as moderator showed that daily activity started earlier (mean estimate [95% CI] = -1.494 [-1.902, -1.087]) and ended later (mean estimate [95% CI] = 0.667 [0.247, 1.088]) in birds exposed to light pollution. We also found that ALAN was associated with higher metabolic rate (mean estimate [95% CI] = 0.476 [0.037, 0.915]), accelerated reproductive maturation (mean estimate [95% CI] = 0.653 [0.037, 1.268]), and higher nocturnal activity (mean estimate [95% CI] = 0.652 [0.107, 1.198]), and foraging effort (mean estimate [95% CI] = 1.289 [0.650, 1.928]). In contrast, ALAN was associated with disrupted sleep regulation (mean estimate [95% CI] = - 0.491 [-0.963, -0.018]). A non-significant trend also indicated that ALAN may increase the level of activity (mean estimate [95% CI] = 0.442 [-0.060, 0.945]), and no significant effects were detected for the remaining traits (full model outputs are shown in Table 3; Fig. 2).

**Figure 2.**
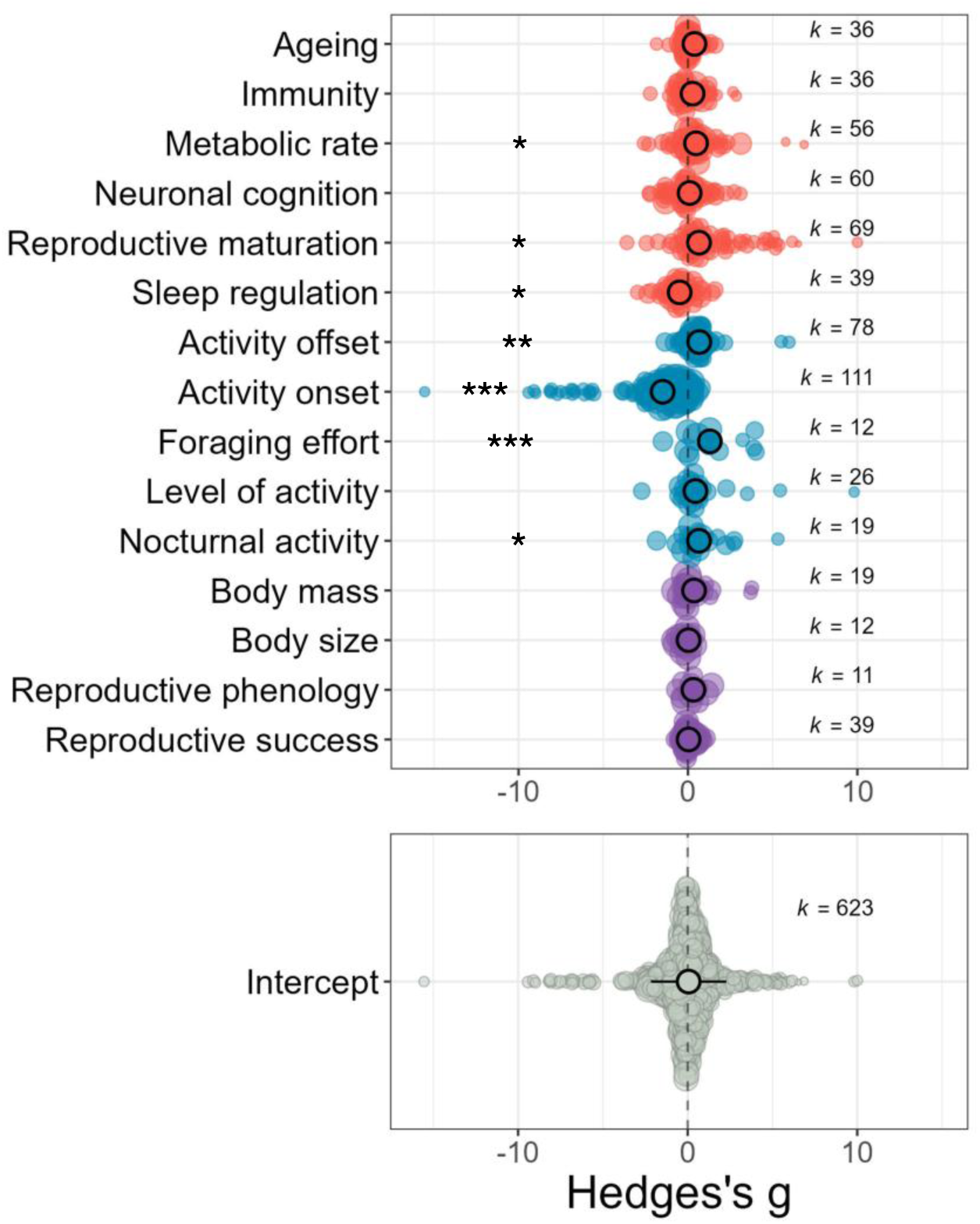
Artificial light at night (ALAN) induced changes in specific physiological and behavioural functional traits. Results from the meta-analytic model including a moderator of different functional traits (y-axis) underpinning physiology (red dots), behaviour (blue dots) and life history (purple dots) traits integrating avian performance. The figure below shows the mean effect size of ALAN across all functional traits fitted by a phylogenetic multi-level (intercept only) meta-analytic model. Both orchard plots show estimates for Hedges’s g (x-axis) along with its 95% confidence intervals (CIs, thick whisker) and 95% prediction intervals (thin whisker). Positive and negative estimates indicate positive and negative effects of ALAN, respectively. The asterisks show the significance level (*P < 0.05, **P < 0.01 and ***P < 0.001) and k represents the number of effect sizes for each level. Outputs of full models 1 and 2 are shown in Table 3.

**Table 3.** Summary of the meta-analytical models on the effects of light pollution on avian functional traits. Intercept only model displays the entire model summary with the associated heterogeneity (I²) explained for each of the random factors, I² total (total heterogeneity), I² ACC (study observation), I² sID (individual effect size observation), I² species ID (species), and I² phylogeny (phylogenetic correlation between species). The global model included avian functional traits as moderator (15 levels, k = 623). Bold estimates indicate confidence intervals (CIs) that did not overlap zero. For each level, k (number of effect sizes), n (number of studies), and species (number of species) is reported.

| Model 1 |  | Mean estimate [CI95%] | k | n | species |
| --- | --- | --- | --- | --- | --- |
| Intercept only model<br>$I^2$ total= 89.90<br>$I^2$ ACC= 16.51<br>$I^2$ sID= 57.60<br>$I^2$ speciesID = 1.79<br>$I^2$ Phylogeny= 14.01 | | 0.056 [-0.482, 0.593] | 623 | 36 | 30 |
| **Model 2** Avian functional traits |  |  |  |  |  |
| Physiology | Ageing | 0.385 [-0.088 , 0.857] | 36 | 6 | 3 |
|  | Immunity | 0.272 [-0.222 , 0.766] | 36 | 5 | 3 |
|  | **Metabolic rate** | **0.476 [0.037 , 0.915]** | **58** | **9** | **5** |
|  | Neuronal cognition | 0.099 [-0.037 , 0.915] | 60 | 5 | 3 |
|  | **Reproductive maturation** | **0.653 [0.037 , 1.268]** | **69** | **3** | **3** |
|  | **Sleep regulation** | **-0.491 [-0.963 , -0.018]** | **39** | **9** | **4** |
| Behaviour | **Activity offset** | **0.667 [0.247 , 1.088]** | **78** | **8** | **8** |
|  | **Activity onset** | **-1.494 [-1.902 , -1.087]** | **111** | **12** | **14** |
|  | **Foraging effort** | **1.289 [0.650 , 1.928]** | **12** | **3** | **8** |
|  | Level of activity | 0.442 [-0.060 , 0.945] | 26 | 6 | 3 |
|  | **Nocturnal activity** | **0.652 [0.107 , 1.198]** | **19** | **4** | **6** |
| Life history | Body mass | 0.361 [-0.145 , 0.866] | 19 | 13 | 6 |
|  | Body size | 0.033 [-0.527 , 0.593] | 12 | 6 | 3 |
|  | Reproductive phenology | 0.323 [-0.289 , 0.935] | 11 | 5 | 5 |
|  | Reproductive success | 0.037 [-0.416 , 0.491] | 39 | 9 | 5 |

### Influence of intra-specific variation

When analysing adults and nestlings separately, we found that ALAN-induced changes in functional traits in adults but not in nestlings (Model 4-5; Table 4). Adults exposed to ALAN showed increased body mass (mean estimate [95% CI] = 1.256 [0.580, 1.932]), accelerated ageing (mean estimate [95% CI] = 0.568 [0.017, 1.12]), and disrupted sleep regulation (mean estimate [95% CI] = -0.476 [-0.879, -0.072]). Models assessing sex-specific effects (Models 6-7; Table 4) showed that ALAN advanced activity onset in males (mean estimate [95% CI] = - 2.393 [-3.831, -0.955]), but not in females (mean estimate [95% CI] = 0.096 [–0.547, 0.740]). Conversely, ALAN induced reproductive maturation (mean estimate [95% CI] = 0.881 [0.225, 1.536]), enhanced neuronal cognition (mean estimate [95% CI] = 1.111 [0.236, 1.986]), and increased foraging effort (mean estimate [95% CI] = 1.235 [0.397, 2.073]) in females, with no comparable changes in males. Full models outputs comparing traits between life-stages and sexes are shown in Table 4; Fig. 3A-B.

**Figure 3.**
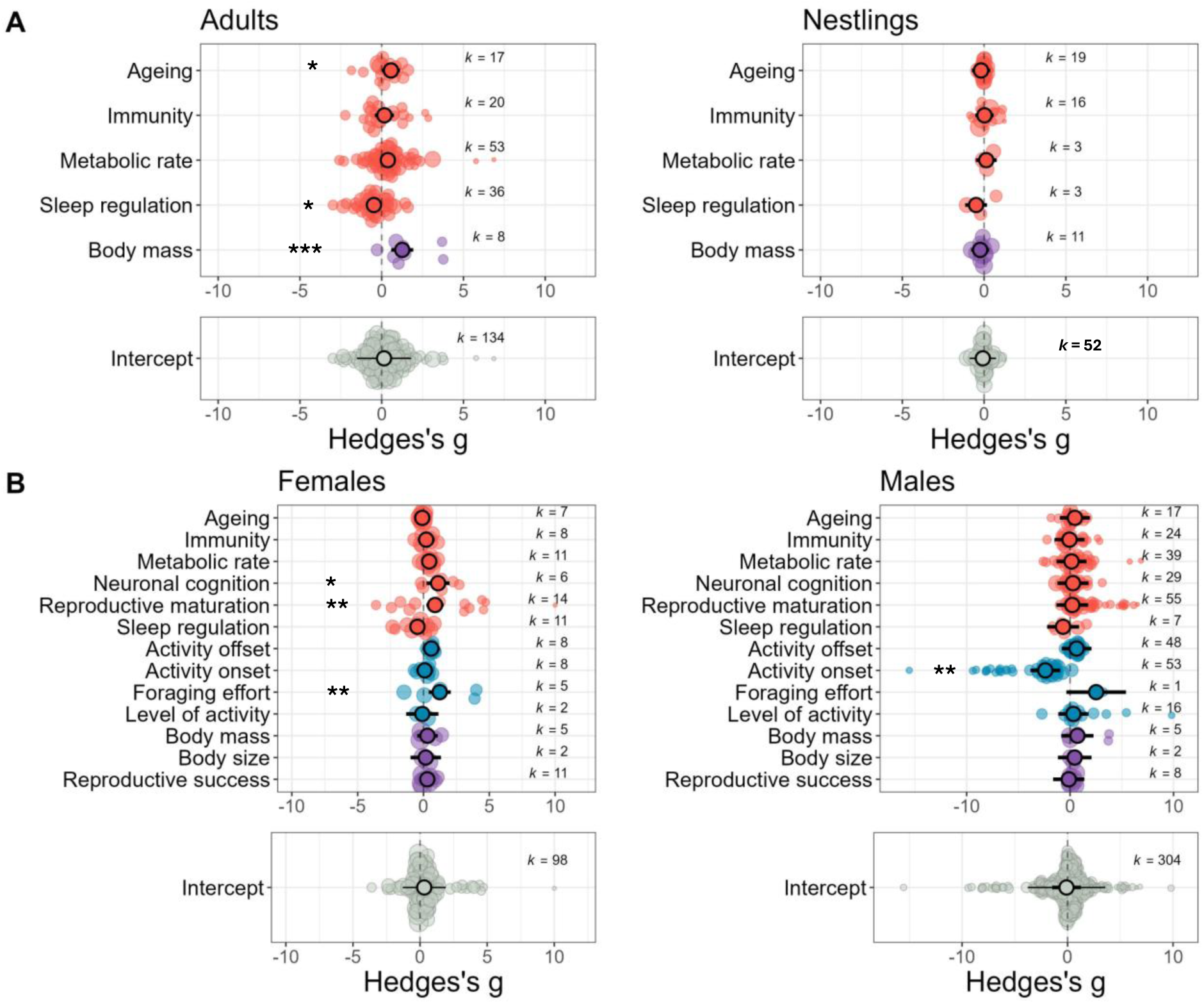
Artificial light at night (ALAN) induced changes on specific functional traits only in adults and females. Results from meta-analytic models comparing the effects of ALAN between A) life-stages (adults vs nestlings) and B) sexes (females vs males). Avian functional underpinning physiology (red dots), behaviour (blue dots) and life histories (purple dots) that integrate avian performance are shown in y-axis. The plots below show the mean effect size of ALAN across all functional traits. The orchard plots show estimates for Hedges’ g along with its 95% confidence intervals (CIs, thick whisker) and 95% prediction intervals (thin whisker) in x-axis. Positive and negative estimates indicate positive and negative effects of ALAN, respectively. The asterisks show the significance level (*P < 0.05, **P < 0.01 and ***P < 0.001) and k represents the number of effect sizes for each level. Outputs of full models 4–7 are shown in Table 4.

**Table 4.** Influence of intra-specific variation on the effects of light pollution on avian functional traits. We compared effects between life-stage (adults vs nestlings) by running separate subset models for adults and nestlings. To enable direct comparisons, subsets included only functional traits reported in both life-stages. The same approach was applied to compare effects between sexes (females vs males). The table shows the results of each meta-analytic model with bold estimates indicating confidence intervals (CIs) that did not overlap zero. For each level, *k* represents the number of effect sizes and n the number of studies.

| Functional trait | Estimate [CI95%] | k | n | Estimate [CI95%] | k | n |
| --- | --- | --- | --- | --- | --- | --- |
|  | **Adults**<br>**Model 4** |  |  | **Nestlings**<br>**Model 5** |  |  |
| **Ageing** | **0.568 [0.017 , 1.12]** | **17** | **3** | -0.187 [-0.750 , 0.375] | 19 | 3 |
| Immunity | 0.167 [-0.416 , 0.749] | 20 | 2 | 0.023 [-0.539 , 0.585] | 16 | 3 |
| Metabolic rate | 0.378 [-0.006 , 0.762] | 53 | 7 | 0.133 [-0.506 , 0.772] | 3 | 2 |
| **Sleep regulation** | **-0.476 [-0.879 , -0.072]** | **36** | **7** | -0.488 [-0.156 , 0.180] | 3 | 2 |
| **Body mass** | **1.256 [0.58 , 1.932]** | **8** | **5** | -0.240 [-0.794 , 0.313] | 11 | 8 |
|  | **Females**<br>**Model 6** |  |  | **Males**<br>**Model 7** |  |  |
| Ageing | -0.092 [-0.711 , 0.526] | 7 | 3 | 0.465 [-0.996 , 1.927] | 17 | 3 |
| Immunity | 0.207 [-0.435 , 0.849] | 8 | 2 | -0.054 [-1.504 , 1.395] | 24 | 2 |
| Metabolic rate | 0.437 [-0.084 , 0.958] | 11 | 2 | 0.144 [-1.302 , 1.59] | 39 | 4 |
| Neuronal cognition | **1.111 [0.236 , 1.986]** | **6** | **1** | 0.28 [-1.226 , 1.785] | 29 | 2 |
| Reproductive maturation | **0.881 [0.225 , 1.536]** | **14** | **1** | 0.214 [-1.319 , 1.747] | 55 | 2 |
| Sleep regulation | -0.458 [-1.001 , 0.085] | 11 | 3 | -0.679 [-2.224 , 0.865] | 7 | 3 |
| Activity offset | 0.591 [-0.066 , 1.248] | 8 | 1 | 0.63 [-0.805 , 2.065] | 48 | 1 |
| Activity onset | 0.096 [-0.547 , 0.74] | 8 | 1 | **-2.393 [-3.831 , -0.955]** | **53** | **2** |
| Foraging effort | **1.235 [0.397 , 2.073]** | **5** | **1** | 2.519 [-0.363 , 5.401] | 1 | 1 |
| Level of activity | -0.087 [-1.313 , 1.139] | 2 | 1 | 0.325 [-1.139 , 1.789] | 16 | 2 |
| Body mass | 0.305 [-0.475 , 1.086] | 5 | 4 | 0.712 [-0.836 , 2.26] | 5 | 4 |
| Body size | 0.174 [-0.982 , 1.331] | 2 | 2 | 0.446 [-1.177 , 2.069] | 2 | 2 |
| Reproductive success | 0.292 [-0.238 , 0.821] | 11 | 3 | -0.128 [-1.638 , 1.382] | 8 | 3 |

### Influence of inter-species traits

We found that ALAN particularly affected fully migratory species (Model 8; mean estimate [95% CI] = 0.081 [0.357, 1.805]), while the effects of ALAN were not significant for partial migrants (mean estimate [95% CI] = 0.873 [-0.006, 1.751]), and resident species (mean estimate [95% CI] = 0.725 [-0.179, 1.629]) (Fig. 4A). Although habitat density had no significant influence on light pollution impact (Model 8; Fig. 4B), there was a tendency for ALAN effects to be negative in species from open habitats (mean [95% CI] = –0.709 [–1.464, 0.046]), while no effects were detected in those from semi-open or dense habitats. In Model 8, we used Type II tests to evaluate each moderator’s significance while accounting for moderate collinearity among them (GVIFs < 5) All moderators significantly contributed to heterogeneity in effect sizes: migration (QM = 8.78, df = 3, *p =* 0.032), habitat density (QM = 229.64, df = 3, *p* < 0.0001), and functional trait (QM = 54.44, df = 13, *p* < 0.0001). Model 9 for study environment (captive vs. wild; Fig. 4C) and Model 10 for population trend (decreasing, stable, increasing; Fig. 4D) did not explain variation in ALAN effects. A summary of all models for inter-specific traits is provided in the supplementary material, Table S3.

**Figure 4.**
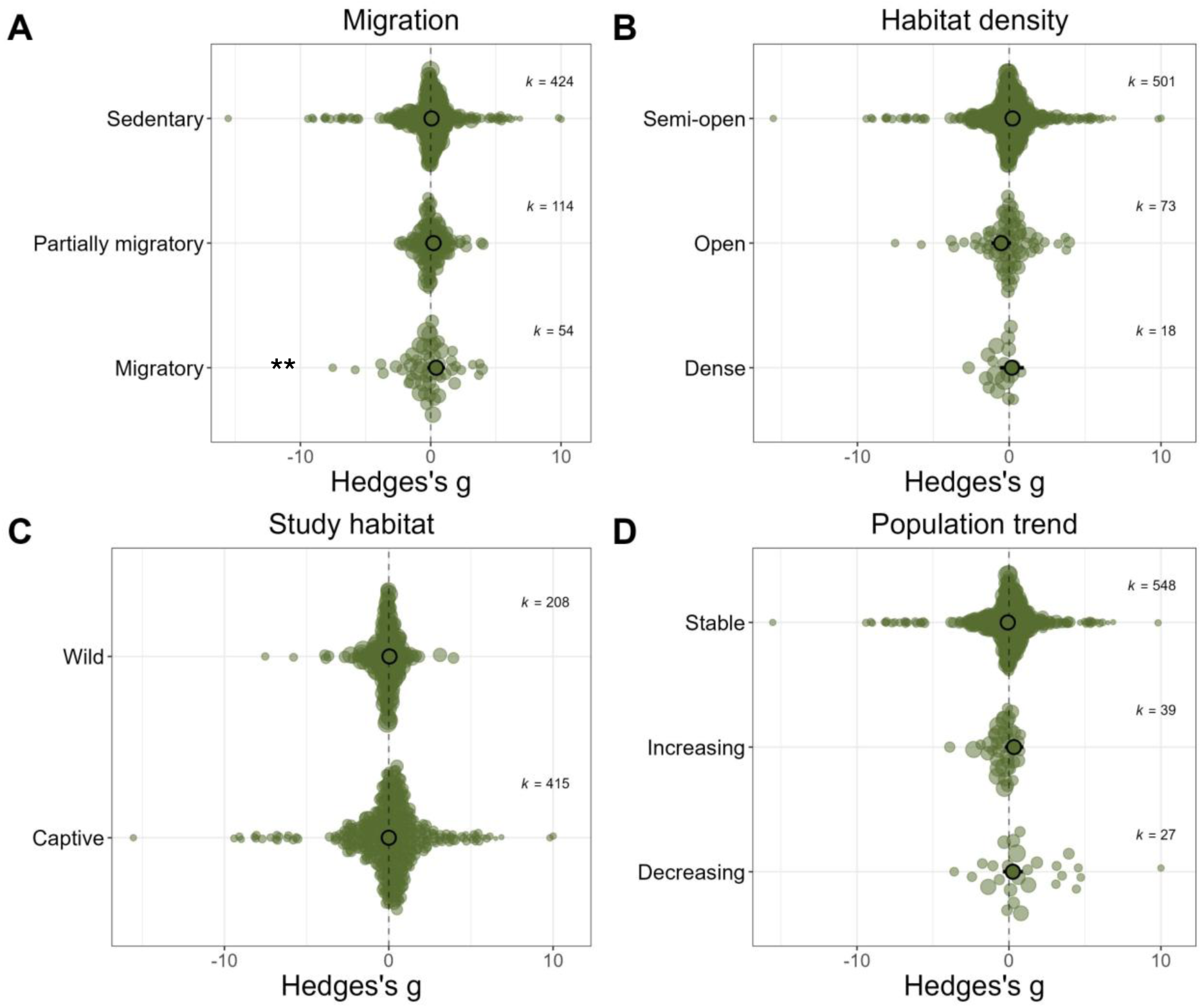
The effects of artificial light at night (ALAN) appear stronger on avian functional traits of migratory species, while other inter-specific traits have no significant influence. Results from the meta-analytic models including A) migration, B) habitat density, C) study habitat and D) population trait as moderators along with avian functional traits. The orchard plots show model estimates for Hedges’s g (x-axis) along with its 95% Confidence Intervals (CIs, whiskers), and the categories of each inter-specific trait in y-axis. Positive and negative estimates assume a positive or negative effect of ALAN, respectively. The asterisks show the significance level (**P < 0.01) and k represents the number of effect sizes for each level. Outputs of full models 8-10 are shown in Table S3.

### Subset analysis on circadian clock and experimental regimes

Model 11 for circadian clock functioning showed significant overall negative effects of ALAN. These effects were significant only if measured at night, and light intensity moderated the magnitude of these effects (details provided in supplementary material, Table S5; Fig. S2A-C). We did not find significant effects of light intensity on the overall functional traits (Model 12; Table 6). However, specific-trait models revealed that higher light intensity was linked to accelerated ageing, reduced sleep drive, earlier activity onset, and increased nocturnal activity (Fig. S3B, G, I, K). Longer ALAN exposure correlated with enhanced reproductive success (Model 13; Fig. S4C). Positive effects of ALAN on neuronal cognition occurred only during the day, whereas ALAN-induced reproductive maturation was independent of sampling time (Model 14; Fig. S5D-E). Notably, increased foraging effort and sleep disruption, previously observed in the global model, were significant only at night (Model 14; Fig. S5F-G). ALAN accelerated ageing in the liver but not in the hypothalamus, and reduced immunity was observed in the hypothalamus but not in the spleen (Model 15; Fig. S6A-B). Detailed model outputs on experimental regimes are shown in supplementary material Table S6.

### Publication bias

The dataset showed evidence of small-study effects, indicating that studies with smaller sample sizes tend to report larger and more significant effects than those with larger samples (mean estimate [95% CI] = 1.025 [0.187, 1.864]; supplementary material, Fig. S7A), though this effect only explained a low proportion of the variance (R^2^_marginal_= 0.023). Following Nakagawa et al., (2022), the intercept from this small-study effects model provided an unbiased estimate of the overall effect size (intercept mean estimate [95% CI] = -0.315 [-1.011, 0.380]). This unbiased estimate was lower than the intercept-only model estimate (Model 1; mean estimate [95% CI] = 0.056 [-0.482, 0.593]), suggesting an inflation of the naïve estimate towards positive values. However, both estimates overlapped considerable and included zero indicating that while small-study effects were detected, they did not substantially alter the overall conclusions. Conversely, we did not find evidence of time lag bias (mean estimate [95% CI] = 0.054 [-0.006, 0.113]; supplementary material, Fig. S7B), the model showed limited proportion of variance explained by time lag bias (R^2^_marginal_= 0.029).

## DISCUSSION

This meta-analysis revealed consistent shifts in physiological and behavioural functional traits under ALAN across birds. In order of effect magnitude: ALAN most strongly advanced daily activity onset, followed by increased foraging effort, delayed daily activity offset, accelerated reproductive maturation, elevated levels of nocturnal activity, disrupted sleep regulation and increased metabolic rate. Exposure to ALAN was associated with increased body mass, accelerated ageing and reduced sleep regulation in adults, but not in nestlings. While females exposed to ALAN showed increased foraging effort, improved neuronal cognition and accelerated reproductive maturation, males largely advanced their daily activity onset. Interestingly, light pollution had stronger effects on the performance of migratory birds, compared to partial migrants and sedentary species. These effects were phylogenetically heterogeneous but consistent across study environments, habitat openness, and species’ extinction risk. Remarkably, our study adds to previous meta-analyses showing that higher light intensities amplified these disruptions, underscoring the need for policies limiting maximum illumination, as light levels below 2 lux caused significantly less disturbance.

The intercept-only model showed no overall effect of ALAN, which is not surprising given that functional traits influence avian performance in opposing directions (e.g. advancing daily activity onset, delaying activity offset). High heterogeneity in Hedges’s g indicates that ALAN effects are trait- and context-dependent (Fig. 2), with most variance arising from within-study variation and phylogenetic relationships among species. Looking across all functional traits, ALAN most strongly affected the timing of activity, with birds advancing their onset into the early morning and extending their offset into the evening, indicating a broad temporal expansion of activity. This behavioural shift was followed by increased foraging effort, particularly at night (supplementary material, Fig. S5G), and by elevated nocturnal activity, consistent with partial behavioural adaptation to illuminated conditions. Earlier onset and extended daily activity are among the most frequently reported behavioural responses to light pollution (Da Silva *et al*. 2016; Da Silva & Kempenaers 2017; Dominoni *et al*. 2014; Pease & Gilbert 2025). We found that the earlier start of activity under ALAN was specific to males (Table 4; Fig. 3B). Early activity onset is thought to provide potential benefits for male singing and extra-pair opportunities (Dominoni & Nelson, 2018; Murphy et al., 2008), suggesting that ALAN may amplify traits already under selection. However, reproductive success also depends on female availability and reproductive strategies (Brouwer & Griffith, 2019; Westneat & Stewart, 2003). Recent work has revealed that urbanisation and ALAN can also advance activity timing in females (Capilla-Lasheras *et al*. 2025; Champenois *et al*. 2025; McGlade *et al*. 2023; Womack *et al*. 2023). This indicates that previous studies were likely male-focused and underscores the need to include female responses in future research.

Beyond sex-specific responses, ALAN disrupts circadian rhythms through changes in core clock gene expression and melatonin suppression, which can both shift daytime activity and promote increased nocturnal activity (Brandstätter 2002; Cassone 2014; Dominoni *et al*. 2022). Additionally, as ALAN improves nocturnal vision, it may stimulate birds to forage longer into the night, but potentially increasing interspecific competition and predation risk (Bonter *et al*. 2013; Leveau 2020). Sleep disruption is another trait possibly linked to increased nocturnal activity. We found that ALAN disrupted avian sleep regulation (Fig. 2), particularly at night (supplementary material, Fig. S5F). This effect is not surprising in diurnal species and can connect both physiological and behavioural responses to ALAN (Table 3; Fig. 2) through nocturnal melatonin suppression (Bentley 2001; Cassone 1990; Grubisic *et al*. 2019). Melatonin, central to initiating and maintaining sleep, was consistently suppressed under ALAN in our study. This aligns with extensive evidence reported across vertebrates (Cassone & Kumar 2022; Grubisic *et al*. 2019; Grunst & Grunst 2023) including recent meta-analytical evidence (Yang *et al*. 2024). We also observed a dysregulation of molecular modulators of sleep onset (e.g., *sik3* mRNA expression), and wakefulness promotion (e.g., *camkII* mRNA expression and oxalate). These modulators are also interconnected with metabolic processes (Batra *et al*. 2020; He *et al*. 2023; Li *et al*. 2023), likely reinforcing the link between disrupted circadian rhythms, poor sleep, and altered energy balance. Importantly, this overall expansion of activity likely incurred physiological costs, as increased locomotor effort elevates energy demands and metabolic rate (Batra *et al*. 2019), a pattern supported by our finding of higher metabolic rate (Table 3; Fig. 2). Metabolic rate is a functional trait interconnected not only with activity levels but also with elevated corticosterone release as a co-effect of ALAN-induced suppression of nocturnal melatonin (Jimeno & Verhulst 2023; Michael Romero 2002; Sapolsky *et al*. 2000). Finally, we observed accelerated reproductive maturation under ALAN, particularly in endocrine markers that trigger breeding. While this trait falls outside the immediate behavioural-metabolic pathway, it is functionally tied to the circadian system and melatonin-mediated photoperiodic regulation (Helm *et al*. 2024; Helm & Liedvogel 2024). We observed that ALAN induced reproductive maturation through the release of reproductive hormones and the expression of reproduction-related genes (supplementary material, Table 3; Fig. 2). These effects were specific to females (Table 4; Fig. 3B), with no significant effects detected in males. Female reproductive physiology is likely more sensitive to environmental cues (e.g., photoperiod, stress and temperature), as reproductive timing and hormonal cycles are tightly linked to circadian and seasonal regulation (Helm *et al*. 2017). Such female-related effects could also arise because males are generally less sensitive to physiological changes in their reproductive activation (Ball & Ketterson 2007; McEwen & Wingfield 2003). Thus, elevated reproductive readiness under ALAN may carry long-term fitness consequences, including mismatches with optimal breeding conditions and increased energetic demands on females. Notably, only females exhibited enhanced neuronal cognition and foraging effort under ALAN (Table 4; Fig. 3B). This sex-specific sensitivity may reflect females’ greater physiological and behavioural investment during reproduction, particularly in egg formation, incubation, and provisioning (Love *et al*. 2005; Vézina & Salvante 2010). Consequently, females are likely more responsive to photoperiodic cues like ALAN that alter timing and energy allocation.

Surprisingly, we did not detect any overall effects of ALAN on immune traits. This may be because such effects often emerge in combination with additional stressors, such as pathogen exposure or immune suppressants (Markowska *et al*. 2017; Tognini *et al*. 2018; Ziegler *et al*. 2021), which were not consistently reported across our dataset. Recent evidence also indicates that ALAN effects on avian immunity are tissue-dependent (Dominoni *et al*. 2022), a finding that aligns with the tissue-dependent patterns observed in our subset analysis (supplementary material, Table S6; Fig. S6B). Additionally, several studies assessed immune traits across different tissues and light intensities using small sample sizes, and this may have introduced small-study effects (Fig S7) and contributed to the limited evidence for ALAN’s impact on immunity.

Our analysis showed life stage-specific differences in avian response to ALAN. Adults displayed accelerated ageing, particularly in terms of the expression of ageing-related genes (Supplementary material, Table S1). The expression of genes encoding key antioxidant enzymes (*gst, sod3, cat1, sirt1*) and promoting mitochondrial biogenesis (*IGF1*) is remarkably sensitive to ALAN due to their tight connection to the circadian clock (Batra *et al*. 2022; Dominoni *et al*. 2022; Taufique *et al*. 2018), though whether altered expression of these genes translates to accelerated organismal ageing and shorter lifespan remains unknown. Adult birds additionally showed increased body mass under ALAN (Table 4; Fig. 3A), which may result from greater energy intake through extended foraging under ALAN (Da Silva *et al*. 2016; de Jong *et al*. 2016; Titulaer *et al*. 2012) and disrupted endocrine hunger regulation (Mahdavi *et al*. 2025). The absence of sleep disruption in nestlings, compared to clear ALAN effects in adults, suggests that age-related vulnerabilities to ALAN emerge later in life, once circadian, endocrine, and behavioural systems are fully developed. However, this also raises the question of how circadian endocrine rhythms emerge during early life and the role of ontogeny, both of which remain poorly understood despite evidence of circadian activity before hatching (Wellard *et al*. 2025). Hence, ALAN may act as an age-dependent stressor, with stronger consequences for adult maintenance and senescence pathways.

On the inter-specific level, we confirmed that the overall effects of ALAN on the functional traits underpinning avian performance are significantly higher in long-distance migrants compared to partial migrants and sedentary species (Table S3; Fig. 4A). Migratory species rely on photoperiodic cues to regulate migratory timing and physiological preparation, making them likely more sensitive to ALAN (Delmore *et al*. 2020; Helm & Liedvogel 2024; Rowan 1930). They also cross landscapes with varying light levels of ALAN during nocturnal movements, which disrupts their natural light cues and have been shown to alter their navigation (Burt *et al*. 2023; Cabrera-Cruz *et al*. 2018). Migrant birds are also known to be attracted to ALAN, which has been particularly detrimental for nocturnal migrants, who account for the higher proportion (∼60%) of collisions with illuminated buildings (Lao *et al*. 2020; Loss *et al*. 2019; Uribe-Morfín *et al*. 2021). Hence, the pronounced physiological and behavioural responses of migrants to ALAN make them a priority for conservation measures and ecological monitoring. By contrast, habitat openness, study environment (e.g., laboratory vs. wild), and species’ population trends (increasing, decreasing, or stable) did not significantly influence ALAN responses (Fig. 4 B-D). This suggests that the physiological and behavioural effects of ALAN are broadly consistent across ecological contexts and population vulnerabilities (Bhagarathi *et al*. 2024; Julliard *et al*. 2004; Mu *et al*. 2021) likely because most data come from stable populations, as research on vulnerable species is often limited. Our dataset, while taxonomically broad, included only 30 avian species (mostly passerines), restricting our ability to capture the full ecological diversity of bird populations.

Remarkably, our subset analysis showed that higher light intensities, in particular exceeding 2 lux, were associated with enhanced nocturnal activity, advanced daily activity onset, accelerated ageing and reduced sleep regulation (supplementary material, Table S6; Fig. S3 B, G, I and K). These amplified effects likely result from increased disruption of melatonin production and circadian gene expression. Brighter light more powerfully suppresses the nocturnal signals that regulate sleep, metabolic rate and cellular repair, which are key processes involved in ageing and daily behavioural rhythms (Dominoni *et al*. 2022; de Jong *et al*. 2016; Moaraf *et al*. 2020a, b). We also found that the number of days since ALAN exposure influenced reproductive success, with longer intervals leading to improved outcomes. This may reflect partial physiological recovery or return to homeostasis from circadian disruption (Helm *et al*. 2024). Additionally, the effects of ALAN on increased foraging effort and sleep disruption were only evident when measured at night, with no difference during the day (supplementary material, Fig. S5F-G). Improvements in neuronal cognition were observed only during the day (supplementary material, Fig. S5D), which is expected given the negative impact of disrupted sleep on nocturnal brain function (Grubisic *et al*. 2019; Moaraf *et al*. 2020b). Interestingly, reproductive maturation was consistently upregulated both during the day and night (supplementary material, Fig. S5E), possibly because reproductive processes are governed by longer-term hormonal changes and do not adjust as rapidly to daily fluctuations (Dominoni *et al*. 2013).

Notably, circadian clock functional traits were not included in our global analysis due to insufficient literature meeting our initial criteria. Yet, our subset analysis revealed consistent ALAN-induced shifts in circadian clock gene expression (e.g., *Per2*, *Cry 1* and *4*, *ck1E* and *Bmal1*), particularly at night when natural dark cycles are disrupted (supplementary material, Table S5; Fig. S2). This disruption of the circadian clock functioning with ALAN may underlie the other physiological and behavioural responses observed in this study. We found most recent studies supporting the robustness of our results. For instance, ALAN induced *Bmal1* phase advancement, leading to earlier activity onset in free-living house sparrows (*Passer domesticus*) and captive zebra finches (*Taeniopygia guttata*) (Coker *et al*. 2025; Hui *et al*. 2025). These observations were also discussed by recent reviews (Grunst & Grunst 2023; Helm & Liedvogel 2024; Wellard *et al*. 2025). Together, these results highlight the need for expanded research on circadian clock gene dynamics, and hormonal regulation, especially in wild populations, to better understand the mechanisms underlying avian responses to ALAN.

Our results do not support the notion that light pollution constrains avian life-history strategies via functional traits. Specifically, we found no evidence of overall changes in body size, reproductive phenology, or reproductive success. This contrasts with meta-analytical evidence linking other anthropogenic disturbances, such as urbanisation and climate warming, to declines in clutch size, advanced breeding phenology, and reduced sexual signal quality (Capilla-Lasheras *et al*. 2022; Halupka *et al*. 2023; Sepp *et al*. 2018). Light pollution may thus exert weaker selective pressure on avian life-history traits than climate change or urbanisation. Birds appear to compensate for ALAN-induced stress via physiological and behavioural adjustments, prioritising core reproductive and survival functions, potentially through phenotypic plasticity (Hau & Goymann 2015) or evolutionary adaptation (Martin 2004; Sepp *et al*. 2018)

## CONCLUSIONS

Our meta-analysis showed that ALAN affected avian performance through behavioural (extended daily activity and increased foraging effort) and physiological (elevated metabolic rate, sleep disruption and induced reproductive maturation) pathways. This aligns with previous evidence that anthropogenic perturbations like ALAN disrupt internal homeostasis, shifting energy acquisition–metabolic expenditure trade-offs with downstream behavioural consequences. In contrast, life-history traits related to growth, reproductive success and survival appeared largely unaffected. This pattern helps explain the paradox often observed in light-pollution research: while ALAN clearly alters physiology and behaviour, its effects on individual fitness and population-level declines are more complex and context-dependent. Birds likely adapt to light-polluted environments, whether through phenotypic plasticity or genetic change: an encouraging indication for future studies of resilience in many species. Importantly, this study underscores the need for policies that protect migratory species and maintain night-time illumination below 2 lux.

## Supporting information

Supplements Meta-ALAN

## ACKNOWLEDGEMENTS

We thank Bart Kempenaers, Frank A. La Sorte, Alexia Mouchet and Marcel Visser for replying to our data queries.

## AUTHOR CONTRIBUTIONS

SD-P and JS conceived the study. SD-P performed the literature search and extracted effect sizes with input from PC-L, DD and JS. PC-L and JS validated effect size extraction. SD-P performed all statistical analysis with substantial advice from PC-L. SD-P wrote the first draft, MC and JS reviewed the first version, and all authors contributed substantially to further revisions of the manuscript.

## DATA ACCESIBILITY

All R scripts and datasets needed to reproduce the analyses presented in this paper are available at: https://doi.org/10.5281/zenodo.17340025

