## Supplements Meta-ALAN for "Artificial light at night consistently impacts avian physiology and behaviour: a meta-analysis"

Keywords: light pollution, avian performance, systematic review, meta-analysis, circadian biology, physiology, activity onset, foraging effort, reproductive output.

**Supplementary text**

*Calculating mean and standard deviation (SD) from estimates*

We extracted 17 estimates and their standard errors (SE) from linear models comparing experimental (ALAN) and control (DARK) groups. These models represented the estimate as the difference between ALAN and DARK, while the intercept corresponded to the mean of the reference group (usually DARK). We calculated the mean for the compared group (ALAN) by adding the estimate to the intercept, but it was subtracted when the estimate was negative. The standard deviation (SD) was derived from the standard errors using the formula: $SD = \sqrt{(SE^2\_intercept + SE^2\_estimate)}$.

*Difference between baseline and final day of exposure*

We found 26 effect sizes reported at two different time points, before (baseline) and after ALAN exposure, for both ALAN and DARK groups. We calculated the difference in mean and standard deviation (SD) between the baseline and the last day of ALAN following (Cooper et al., 2009). We used the formula $D= \bar{Y}1 - \bar{Y}2$ to calculate the difference in means between two independent groups. The SD of this difference was derived by first calculating the variance of each group by squaring their standard deviations, then computing the variance of the difference between the groups using the formula $\sigma 2X{-}Y= \sigma 2X + \sigma 2Y$ and finally taking the square root of this combined variance to obtain the SD of the difference between the independent variables.

**Supplementary tables**

**Table S1.** Description of the effect sizes included in the meta-analysis on the effects of artificial light at night on avian performance. The table shows the effect sizes assigned to 16 avian functional traits underpinning avian performance, sorted by meta-analytical categories: physiology (red), behaviour (blue) and life history traits (violet) . Effect size are presented with a functional note explaining its inclusion to the specific functional trait. To enable interpretation of the meta-analysis results, we assigned a direction to each effect size. If an increase in the effect size was expected to negatively impact the particular avian functional trait, the direction was negative. When the direction was not reported in the original study, effect sizes expected to have a negative impact were multiplied by -1. The table summarizes 675 effect sizes from ALAN–DARK paired comparisons across 36 studies published between 2006 and 2022.

| Level | Functional trait | Included effect sizes | Functional note | Direction |
|---|---|---|---|---|
| Physiology | Ageing | gst, sod3 and cat1 mRNA expression | Activation of antioxidant enzymes | negative: Higher values enhance antioxidant capacity, thereby slowing ageing |
| | | GSH, GSSG and GSH/GSSG | Molecular antioxidants | negative: Higher values enhance antioxidant capacity, thereby slowing ageing |
| | | TBARS and protein carbonyls | Marker of oxidative stress | positive: Higher values enhance oxidative stress, thereby accelerating ageing |
| | | RTL: relative telomere length | Maintain cellular stability and regulates cellular senescence | negative: Higher values enhance cell stability, thereby slowing ageing |
| | | NOx: nitric oxide concentrations | Inhibition of telomerase and telomere shortening | positive: Higher values shorten telomeres, thereby accelerating ageing |
| | | TAC: total antioxidant activity | Non enzymatic antioxidant capacity | negative: Higher values enhance antioxidant capacity, thereby slowing ageing |
| | | GPX, CAT and SOD | Protect cells against oxidative stress | negative: Higher values enhance antioxidant capacity, thereby slowing ageing |

| | | | | |
|---|---|---|---|---|
| | | Sirt1 mRNA expression | Mediates stress response and promotes longevity | negative: Higher values promotes longevity, thereby slowing ageing |
| | | IGF1 mRNA expression | Hormone promoting mitochondrial biogenesis, respiration and ageing | negative: Higher values promotes longevity, thereby slowing ageing |
| | Circadian clock functioning | Per2 mRNA expression | Regulates circadian phase; responds to light signals with peak of expression at early night | negative at night: lower expression at night indicate effective circadian clock regulation |
| | | Cry1 mRNA expression | Inhibits CLOCK-BMAL1 activity stabilizing circadian rhythm with peak in dark phase | negative at night: lower expression at night indicate effective circadian clock regulation |
| | | Cry4 mRNA expression | Light-dependent magnetoreceptor for daily timing (peak in light phase) | negative at night: lower expression at night indicate effective circadian clock regulation |
| | | ck1E mRNA expression | Regulates phosphorylation and degradation of PER proteins with constant expression across light and dark cycles | negative at night: lower expression at night indicate effective circadian clock regulation |
| | | Bmal1 mRNA expression | Key for circadian clock rhythm generation with peak of expression at early night | positive at night: higher expression at night indicate effective circadian clock regulation |
| | Immunity | NOx: nitric oxide concentrations | Modulates inflammatory process and eliminates pathogens | positive: Higher values enhance immunity |
| | | HP- haptoglobin/ % of bacterial killing | Anti-inflammatory mediator | positive: Higher values enhance immunity |
| | | % bacterial E. colli killing | Anti-bacterial infection | positive: Higher values enhance immunity |
| | | cd36 mRNA expression | Involved in lipid metabolism, innate immunity, and regulates inflammation | positive: Higher values enhance immunity |
| | | ly86 mRNA expression | T cell activation and cytokine production | positive: Higher values enhance immunity |
| | | tlr4 mRNA expression | Innate immune response activation | positive: Higher values enhance immunity |
| | | DEE: Daily energy expenditure | Energy expenditure | positive: Higher values indicate increased metabolism |

| | | | | |
|---|---|---|---|---|
| | | Body temperature | Internal thermal state: higher values support physiological stability | positive: Higher values indicate better self-maintenance |
| | Metabolic rate | Corticosterone from plasma and feathers | Induce stress and coordinate energy resources to cope with environmental challenges, it is related with increases in metabolic rate. | positive: Higher values indicate increased metabolism |
| | | foxo 1 and serbp mRNA expression | Regulates the overall cellular glucose and lipid metabolism | positive: Higher values indicate increased metabolism |
| | | npy mRNA expression | Involved in appetite regulation and stress regulation | positive: Higher values indicate increased metabolism |
| | | ppar alpha mRNA expression | Important regulator of lipid metabolic pathways in body tissues | positive: Higher values indicate increased metabolism |
| | | fgf mRNA expression | Stimulates adipogenesis through the MAPK pathway | positive: Higher values indicate increased metabolism |
| | | nrf1- mRNA expression | Mitochondrial activity (transcription factor) | positive: Higher values indicate increased metabolism |
| | | nr3c1 mRNA expression | Glucocorticoid receptor | positive: Higher values indicate increased metabolism |
| | | nr3c2 mRNA expression | Mineralocorticoid receptor | positive: Higher values indicate increased metabolism |
| | Neuronal cognition | Neuronal density ,DCXir, BDNFir, CDX/BDNFir | Neuronal density | positive: Higher values indicate greater neuronal cognition |
| | | Number of cells related with song production | Song production | positive: Higher values indicate greater neuronal cognition |
| | | caspase2, caspase3 and foxo3 mRNA expression | Initiate apoptosis and neuronal death | negative: Higher values indicate lower neuronal cognition |
| | | creb, syngap, syn 2, syn 2a, syn 2b, egrl mRNA expression | Induce neuronal plasticity | positive: Higher values indicate greater neuronal cognition |
| | | Bdnfr, il-1Beta, tnfr1, nr4a2 mRNA expression | Process sensory information to enable cognitive functions | positive: Higher values indicate greater neuronal cognition |
| | | LH: luteinizing hormone | Induce gonadal maturation - initiate reproductive behaviour | positive: Higher values indicate effective reproductive maturation |
| | | Testosterone / Estradiol | Sex differentiation and initiates reproductive behaviour | positive: Higher values indicate effective reproductive maturation |

| | | | |
|---|---|---|---|
| Reproductive maturation | Testis volume, tubule diameter | Indicator or sperm production and reproductive potential | positive: Higher values indicate effective reproductive maturation |
| | THS-B, GnRH-1 and Dio2 mRNA expression | Activate hormones inducing gonadal maturation | positive: Higher values indicate effective reproductive maturation |
| | STRA 8 and SPO11 mRNA expression | Involved in germ cells development regulating meiotic initiation | positive: Higher values indicate effective reproductive maturation |
| | LHR and HSD3B1 mRNA expression | Activate Leydig cells involved in steroid synthesis | positive: Higher values indicate effective reproductive maturation |
| | CLDN11, SOX 9, FSHR and WT1 | Activate Sertoli cells involved in spermatogenesis | positive: Higher values indicate effective reproductive maturation |
| Sleep regulation | Melatonin levels | Hormone inducing sleep onset during in response to darkness | positive: Higher values indicate effective sleep |
| | achm3 mRNA expression | Awake promoter gene- light receptor | negative at night: Higher values indicate defective sleep |
| | % oxalate levels | Biomarker of sleep loss with higher oxalate resulting in better sleeping | positive: Higher values indicate effective sleep |
| | camkii mRNA expression | Promotes neural excitability; higher expression is linked to less sleep | negative at night: Higher values indicate defective sleep |
| | Sik3 mRNA expression | Sleep inducing kinase | positive: Higher values indicate effective sleep |
| Activity offset | Last activity of the day, activity offset, last parental feeding time, offset of dusk singing, cessation of foraging | Timing of activity cessation | positive: higher values indicate later activity offset |
| Activity onset | First activity of the day, chorus onset time, song initiation time, first nest visit, start of the Dawn Chorus, foraging onset. | The time of the first activity , generally measured as minutes to daylight. | negative: the greater the number of minutes, the earlier the activity begins |
| | Singing onset (min past midnight) | The time of first activity is measured as minutes past midnight. | positive: the greater the number of minutes, the later the activity begins |
| Foraging effort | Self-feeding duration | Duration of food intake; longer times suggest higher energy needs. | positive: Higher values indicate better self-maintenance |
| | Food intake | Amount of consumed food; higher intake reflects greater energy demand | positive: Higher values indicate better self-maintenance |

| | | | | |
|---|---|---|---|---|
| Behaviour | | Preys per minute | Feeding efficiency; higher values indicate improved resource acquisition | positive: Higher values indicate better self-maintenance |
| | Nocturnal activity | Awakening time, nocturnal nest leaving time, nocturnal activity duration, awakening latency, nocturnal bouts, sleep latency | Duration or amount of activity during night dark | positive: longer activity duration indicate higher nocturnal activity |
| | | Sleep amount, sleep onset, Total sleep duration, nocturnal activity bout duration, sleeping bouts | Sleeping parameters | negative: longer sleeping activity indicated lower nocturnal activity |
| | Level of activity | Length of active day, nest entry times, activity duration, activity counts/hr, proportion of activity per hour, total activity | Duration or amount of activity during daylight | positive: Higher values reflect increased activity levels |
| | | Latency | Delay on responding to a stimulus | negative: higher values reflect lower activity |
| Life history traits | Body mass | Body mass, weight, mass gain, fat score, growth rate | Influences metabolic rate, energy allocation, and timing of life-history events | positive: Higher values reflects positive life-history traits |
| | Body size | Tarsus, wing, rectrices and bill length | Involved in survival, reproduction, and energy allocation | positive: Higher values reflects positive life-history traits |
| | Reproductive phenology | Incubation length, laying date, nesting period, fledging date | Shape the reproductive timing and success in life-history strategies | positive: longer time associated with increased reproductive success |
| | Reproductive success | Parental feeding rate, fledging success, fledging number, extrapair mates, clutch size, hatching success, brood size, recruitment rate, extra paternity gain, extra pair young | Shape reproductive success and offspring survival | positive: higher values associated with high reproductive success |
| | | Nest predation | Critical factor influencing reproductive success and offspring survival | negative: Higher values indicate lower reproductive success |

**Table S2.** Summary of the inter-species traits included in the meta-analysis. The table shows the scientific names of the 36 species included in our dataset followed by inter-species traits: migration, habitat type (extracted from AVONET database) and population trend (collected from the IUCN Red List. NA indicates missing data for this particular species.

| Scientific name | Migration | Habitat density | Population trend |
|---|---|---|---|
| *Turdus merula* | Sedentary | semi-open | Increasing |
| *Turdus migratorius* | Migratory | open | Stable |
| *Turdus philomelos* | Migratory | dense | Increasing |
| *Catharus ustulatus* | Migratory | dense | Stable |
| *Sialia mexicana* | Partially migratory | open | Increasing |
| *Erithacus rubecula* | Migratory | dense | NA |
| *Ficedula hypoleuca* | Migratory | dense | Decreasing |
| *Mimus polyglottos* | Migratory | semi-open | Stable |
| *Regulus calendula* | Migratory | dense | Increasing |
| *Sitta europaea* | Sedentary | dense | Stable |
| *Junco hyemalis* | Migratory | semi-open | Decreasing |
| *Zonotrichia albicollis* | Migratory | semi-open | Decreasing |
| *Emberiza bruniceps* | Migratory | open | Stable |
| *Setophaga coronata* | NA | NA | Stable |
| *Fringilla coelebs* | Partially migratory | semi-open | Increasing |
| *Passer montanus* | Sedentary | semi-open | Decreasing |
| *Taeniopygia guttata* | Partially migratory | semi-open | Stable |
| *Cyanistes caeruleus* | NA | NA | Stable |
| *Parus major* | Sedentary | semi-open | Stable |
| *Tachycineta bicolor* | Migratory | open | Stable |
| *Progne subis* | Sedentary | semi-open | Stable |
| *Garrulus glandarius* | Partially migratory | dense | Stable |
| *Corvus splendens* | Sedentary | open | Stable |
| *Charadrius hiaticula* | Partially migratory | open | Decreasing |
| *Charadrius alexandrinus* | Partially migratory | open | Decreasing |
| *Recurvirostra americana* | Partially migratory | open | NA |
| *Pluvialis squatarola* | Migratory | open | Decreasing |
| *Tringa totanus* | Migratory | open | NA |
| *Calidris alpina* | Migratory | open | Decreasing |
| *Coturnix chinensis* | NA | NA | Stable |

**Table S3.** Summary of the meta-analytic mixed-effects models showing no significant influence of inter-species traits on the impact of artificial light at night on avian functional traits. The table show estimates and confidence intervals (CIs) for each category of the species-specific trait investigated. The first model included: migration and habitat density. The second and third models included study habitat and population trend as moderators. For each level, *k* represents the number of effect sizes, *n* the number of studies, and species the number of different species included.

| Category | Estimate [CI95%] | k | n | species |
|---|---|---|---|---|
| **Model 8** | **Migration** | | | |
| Migratory | **0.081 [0.357 , 1.805]** | 55 | 11 | 14 |
| Partially migratory | 0.873 [-0.006 , 1.751] | 114 | 10 | 7 |
| Sedentary | 0.725 [-0.179 , 1.629] | 424 | 24 | 6 |
| **Model 8** | **Habitat density** | | | |
| Dense | 0.178 [-0.609 , 0.964] | 18 | 7 | 7 |
| Open | -0.709 [-1.464 , 0.046] | 73 | 6 | 11 |
| Semi-open | 0.043 [-0.719 , 0.805] | 501 | 29 | 9 |
| **Model 9** | **Study environment** | | | |
| Captivity | 0.004 [-0.575 , 0.582] | 415 | 11 | 6 |
| Wild | 0.046 [-0.433, 0.525] | 208 | 25 | 25 |
| **Model 10** | **Population trend** | | | |
| Decreasing | 0.242 [-0.471 , 0.956] | 27 | 4 | 8 |
| Increasing | 0.321 [-0.349 , 0.991] | 39 | 8 | 6 |
| Stable | -0.091 [-0.643 , 0.461] | 548 | 32 | 14 |

**Table S4.** Description of the subset meta-analytical models evaluating the effects of light pollution on circadian clock, and the influence of experimental conditions: light intensities, days since ALAN exposure, time of day, and tissue types. Model indexes contain a series of sequential numbers from 11 to 15 to facilitate the understanding of the methods and results. 'Data' refers to the effect sizes included in each model index and *k* represents the number of effect sizes. 'Moderators' show the categories included as moderators in each model, 'Details' provide a short description per model and 'Output section' show where to find the model output.

| Model index | Evaluating | Data | K= | Moderators | Details | Output section |
|---|---|---|---|---|---|---|
| 11 | Effects of ALAN on circadian clock | Effect sizes of circadian clock gene expression | 52 | a. Intercept only<br>b. Light intensity<br>c. Time of Day | Effects of ALAN on circadian clock functioning | Table S5; Fig. S2 |
| 12 | | All effect sizes associated to a light intensity (lux) measure | 662 | a. Light intensity + functional trait-1 | Effects of light intensity on the overall functional traits | Table S6; Fig. S3A |
| | | Only effect sizes repeated at different light intensities | 336 | b. Light intensity | Light intensity effects (separate models per functional trait) | Table S6; Fig. S3B-L |
| 13 | Influence of varying experimental regimes | Only effect sizes repeated at different days after ALAN exposure (days) | 52 | Time since ALAN exposure | Days since ALAN exposure effects (separate models per functional trait) | Table S6; Fig. S4 |
| 14 | | Only effect sizes repeated during day and at night | 176 | Time of day (day or night) | Day and time effects of ALAN (separate models per functional trait) | Table S6; Fig. S5 |
| 15 | | Only effect sizes reported for different tissues (liver, spleen, hippocampus, hypothalamus and brain areas) | 143 | tissue type | Tissue-related effects of ALAN (separate models per functional trait) | Table S6; Fig. S6 |

**Table S5.** Results on the effects of artificial light at night (ALAN) on avian circadian clock functioning. We tested these effects in a subset including k= 52 effect sizes reported in two different studies. We conducted a phylogenetic multi-level meta-analysis with an intercept-only model, then, we extended this model by including light intensity as a moderator, and we ran an additional model with time of day as a moderator. The results suggest that ALAN significantly affects circadian clock functioning which increases with light intensity, and these effects are significant at night when compared to the day measure. The table shows estimates for each model in separate rows. Bold estimates indicate confidence intervals (CIs) that did not overlap zero.

| **Circadian clock functioning<br>Model 11** | |
|---|---|
| Model | Estimate [CI95%] |
| Intercept only | **-0.540 [-0.814, -0.265]** |
| Light intensity | **-0.114 [-0.226, -0.002]** |
| Time of the day | Day: -0.448 [-0.905, 0.009<br>**Night: -0.596 [-0.945, -0.247]** |

**Table S6**. Summary of the meta-models testing the influence of experimental regimes (light intensity, days since ALAN exposure, time of day and tissue type) on the avian functional traits' responses to artificial light at night. We assessed subsets including effect sizes that were measured at different experimental conditions and ran a model per each functional trait. The effects of light intensity were also assessed in two subsets, one including all the effects sizes from which light intensity was reported and, and another including effect sizes measured at different light intensities. The tissue-type subset included effect sizes measured in eight brain regions associated with learning, all grouped as 'forebrain regions'. The table shows model estimates for each functional trait with bold estimates indicating confidence intervals (CIs) that did not overlap zero. The number of effect sizes (*k)*, studies (*n)* and species are reported for each level.

| **Functional trait** | **Estimate [CI95%]** | **k** | **n** | **species** |
|---|---|---|---|---|
| **Model 12** | **Light intensity (lux)** | | | |
| Overall functional traits | 0.006 [-0.004, 0.016] | 662 | 35 | 30 |
| **Ageing** | **0.203 [0.058, 0.349]** | **12** | **1** | **1** |
| Immunity | -0.072 [-0.197, 0.054] | 18 | 1 | 1 |
| Metabolic rate | -0.039 [-0.186, 0.108] | 34 | 2 | 1 |
| Neuronal cognition | -0.143 [-0.301, 0.015] | 17 | 3 | 2 |
| Reproductive maturation | 0.378 [0.280, 0.475] | 54 | 1 | 1 |
| **Sleep regulation** | **-0.150 [-0.255, -0.044]** | **18** | **2** | **2** |
| Activity offset | -0.010 [-0.078, 0.058] | 55 | 3 | 2 |
| **Activity onset** | **-1.045 [-1.276, -0.814]** | **58** | **3** | **2** |
| Level of activity | 0.155 [-0.196, 0.505] | 16 | 2 | 1 |
| **Nocturnal activity** | **1.168 [0.575, 1.760]** | **4** | **1** | **1** |
| Body mass | 0.877 [-0.246, 2.006] | 2 | 1 | 1 |
| **Model 13** | **Days after ALAN exposure** | | | |
| Immunity | 0.007 [-0.006, 0.019] | 12 | 1 | 1 |
| Reproductive maturation | -0.005 [-0.123, 0.114] | 11 | 1 | 1 |
| **Reproductive success** | **0.092 [0.040, 0.145]** | **8** | **1** | **1** |
| Activity offset | 0.012 [-0.042, 0.018] | 56 | 2 | 1 |
| Activity onset | 0.03 [-0.091, 0.151] | 56 | 2 | 1 |

| Model 14 | Time of sampling | | | |
|---|---|---|---|---|
| Ageing | Day: 0.155 [-0.617 , 0.927] | 7 | 2 | 2 |
| | Night: 0.282 [-0.465 , 1.029] | 7 | 2 | 2 |
| Immunity | Day: -0.187 [-0.802 , 0.427] | 10 | 2 | 2 |
| | Night: 0.032 [-0.51 , 0.575] | 10 | 2 | 2 |
| Metabolic rate | Day: 0.291 [-0.237 , 0.818] | 23 | 5 | 4 |
| | Night: 0.426 [-0.076 , 0.928] | 25 | 5 | 4 |
| **Neuronal cognition** | **Day: 1.932 [0.298 , 3.567]** | **3** | **1** | **1** |
| | Night: -0.025 [-0.883 , 0.833] | 3 | 1 | 1 |
| **Reproductive maturation** | **Day: 0.803 [0.268 , 1.339]** | **24** | **1** | **1** |
| | **Night: 0.574 [0.25 , 0.898]** | **24** | **1** | **1** |
| **Sleep regulation** | Day: 0.117 [-0.271 , 0.506] | 15 | 7 | 4 |
| | **Night: -1.02 [-1.428 , -0.612]** | **15** | **7** | **4** |
| **Foraging effort** | Day: -0.686 [-1.854 , 0.481] | 2 | 1 | 1 |
| | **Night: 3.953 [2.536 , 5.371]** | **2** | **1** | **1** |
| Level of activity | Day: -0.235 [-1.063 , 0.593] | 6 | 4 | 3 |
| | Night: 0.259 [-0.551 , 1.069] | 6 | 4 | 3 |
| **Model 15** | **Tissue type** | | | |
| **Ageing** | Hypothalamus: 0.342 [-0.318 , 1.003] | 6 | 1 | 1 |
| | **Liver: 0.79 [0.237 , 1.342]** | **6** | **1** | **1** |
| **Immunity** | **Hypothalamus: -0.634 [-1.258 , -0.01]** | **6** | **1** | **1** |
| | Spleen: -0.037 [-0.493 , 0.419] | 12 | 1 | 1 |
| Metabolic rate | Hippocampus: -0.304 [-0.878 , 0.269] | 6 | 1 | 1 |
| | Hypothalamus: 0.173 [-0.551 , 0.897] | 10 | 1 | 1 |
| | Liver: -0.418 [-1.262 , 0.426] | 6 | 1 | 1 |
| Neuronal cognition | Forebrain regions: 0.136 [-0.132 , 0.405] | 37 | 4 | 2 |
| | Hypothalamus 0.462 [-0.385 , 1.308] | 6 | 4 | 2 |

**Table S7.** Description of studies excluded during the screening of records following the PRISMA flowchart (details provided in Fig. S1). The research yielded 1,480 records, from which 379 duplicates were removed, resulting in 1,128 unique studies that were screened based on title and abstract screening (phase 1). A full-text screening (phase 2) was performed in 180 records from which 36 studies were included in the meta-analysis. The table lists the reasons for exclusion and the number of studies excluded in each phase.

| Reason for exclusion | Studies excluded in phase 1 | Studies excluded in phase 2 |
|---|---|---|
| 1. Did not retrieve | 11 | 4 |
| 2. Studies outside birds | 86 | - |
| 3. Study on meat/egg production (poultry) and studies on any pedigree bred birds | 472 | 12 |
| 4. The study did not report effects of artificial light at night on a bird in the field or lab | 194 | 49 |
| 5. Unmatching experimental design: Having a control (dark) group exposed to natural light levels at night of max 0.2 lux, and treatment groups with exposure to ALAN up to 100 lx | 141 | 46 |
| 6. Wrong publication type: Data from other sources a patents, videos or vulgarisation science not providing data on means or variance | 23 | 3 |
| 7. Within individual design | - | 10 |
| 8. Light measured from urban areas compared to rural as control groups | - | 5 |
| 9. Experimental design including other stressors (food deprivation, bacterial infection induction etc) | - | - |
| 10. No data available, not provided by authors | | 6 |
| 11. Duplicate | | 9 |
| **Total of excluded studies** | **932** | **135** |

**Supplementary figures**

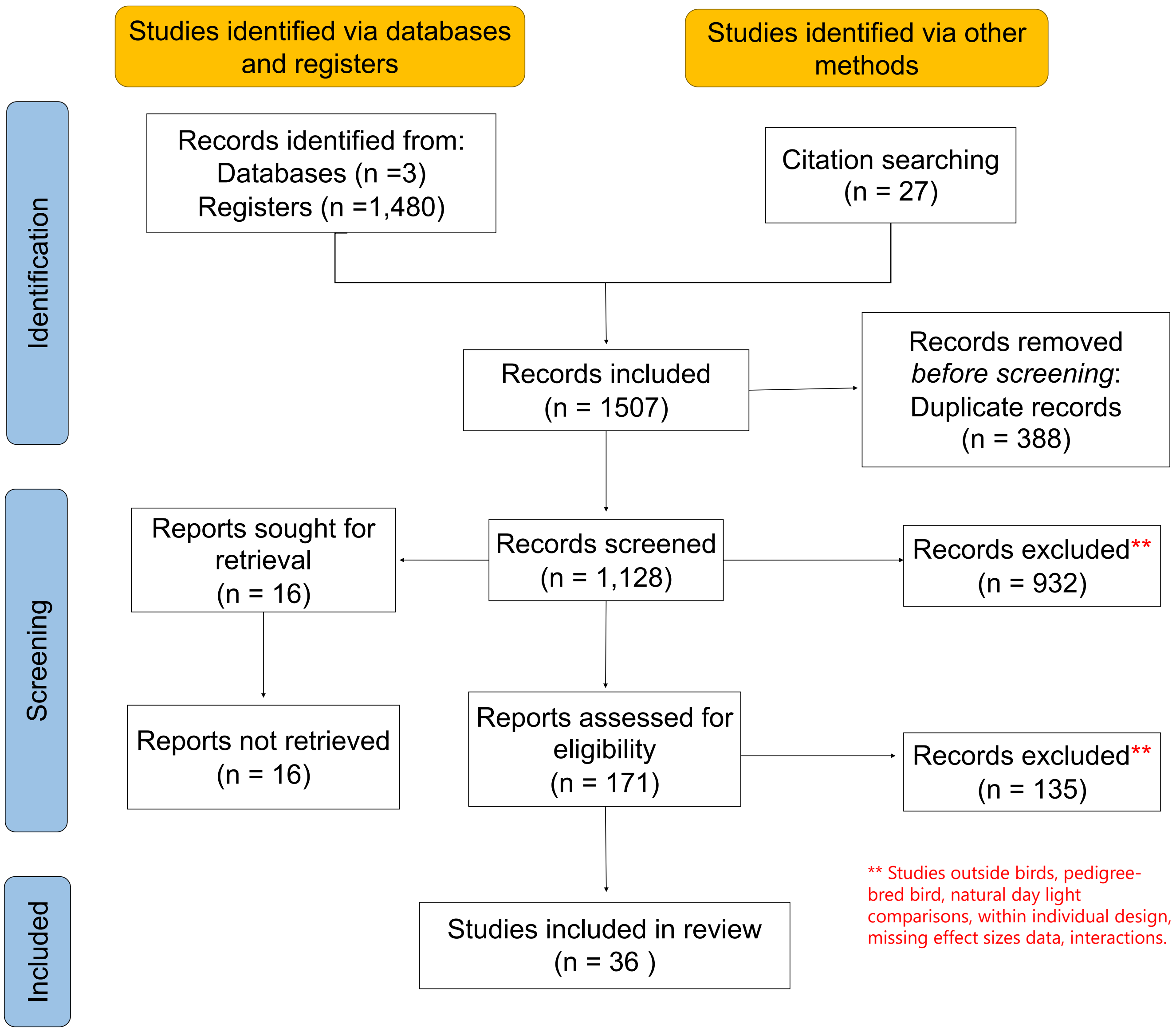


**Figure S1.** Number of studies included in the meta-analysis on the effects of artificial light at night on avian performance. The figure shows the results for each search phase following Preferred Reporting Items for Systematic Reviews and Meta-Analysis (PRISMA) guidelines. The systematic research of published literature was performed using three different databases: Scopus, Web of Science (All databases) and PubMed, and 27 extra studies were identified and included from a previous published meta-analysis on the biological impacts of artificial light at night (Sanders et al., 2020).

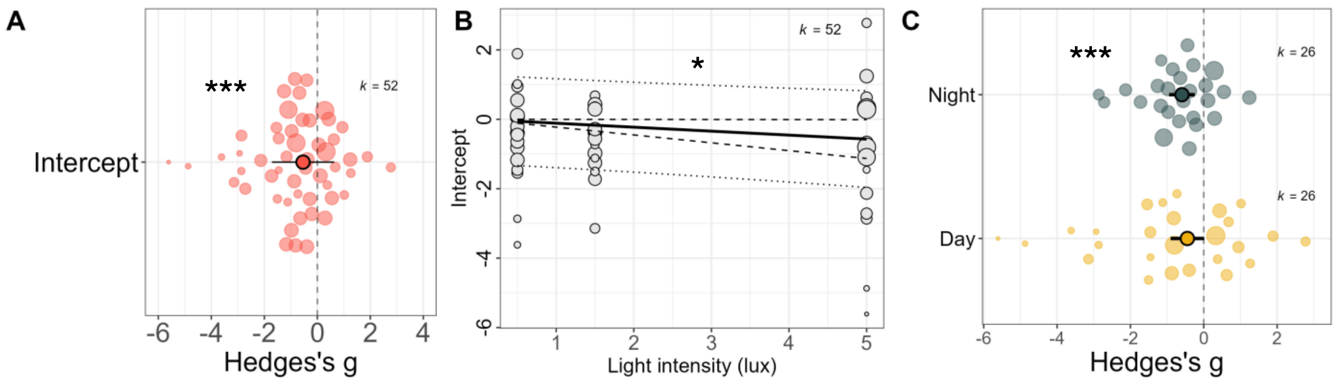


**Figure S2.** Results from the analysis on the effects of artificial light at night on avian circadian clock functioning. A) Orchard plot from the intercept model showed significant and negative effects of ALAN on the overall effect sizes for avian circadian clock. B) Bubble plot showing a decrease in circadian clock functioning associated with increasing light intensity. Hedges' effect sizes are displayed in y-axis and light intensity (measured in lux) in x-axis, the solid line represents the model estimate with the 95% confidence intervals displayed by dashed lines and the 95% prediction intervals by dotted lines. C) Orchard plot for the influence of time of day (day and night) on the impact of ALAN on the avian circadian clock. The orchard plots show the estimates for Hedges's g along with its 95% CIs (thick whisker) and 95% prediction intervals (thin whisker). For all plots, positive and negative estimates indicate positive and negative effects of ALAN, respectively. The asterisks show the significance level (*P < 0.05, **P < 0.01 and ***P < 0.001) and k represents the number of effect sizes for each level. Outputs of full model 11 is shown in Table S5.

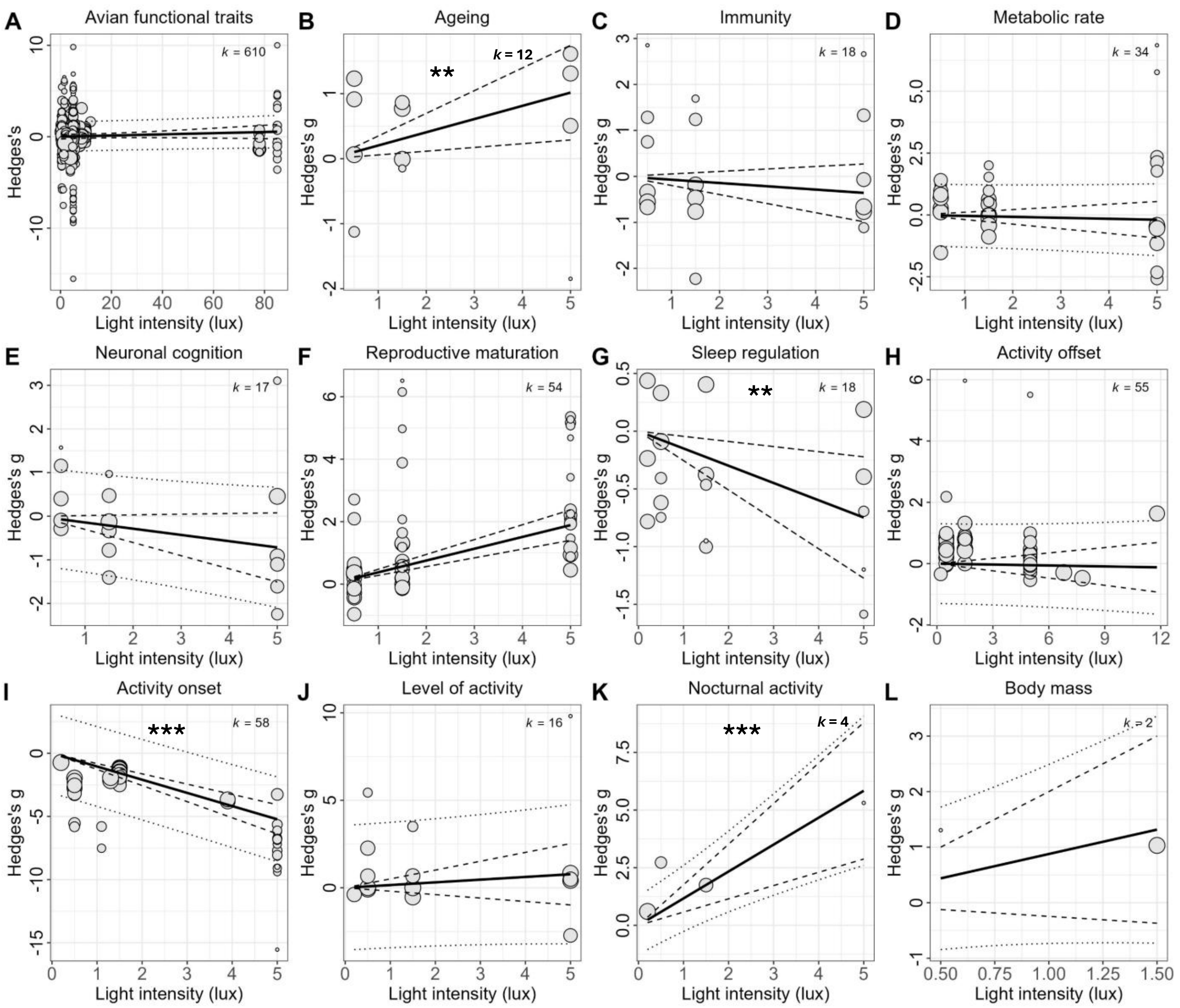


**Figure S3**. Results from the models testing the impact of light intensity on avian function traits. A) Model including all effect sizes from studies that reported a measure of light intensity. Separate models were run from subsets for each functional trait reporting repeated effect sizes at different light intensities: B) ageing, C) immunity, D) metabolic rate, E) neuronal cognition, F) reproductive maturation, G) sleep regulation, H) activity offset, I) activity onset, J) level of activity, K) nocturnal activity and L) body mass. Higher light intensities accelerated ageing and decreased sleep regulation, advanced start of activity and increased nocturnal activity. The bubble plots show Hedges' effect sizes on the y-axis, and light intensity (measured in lux) on the x-axis. The solid line represents the model estimate, with 95% confidence intervals shown as dashed lines and 95% prediction intervals as dotted lines. The asterisks show the significance level (*P < 0.05, **P < 0.01 and ***P < 0.001) and k represents the number of effect sizes for each level. Outputs of full model 12 are shown in Table S6.

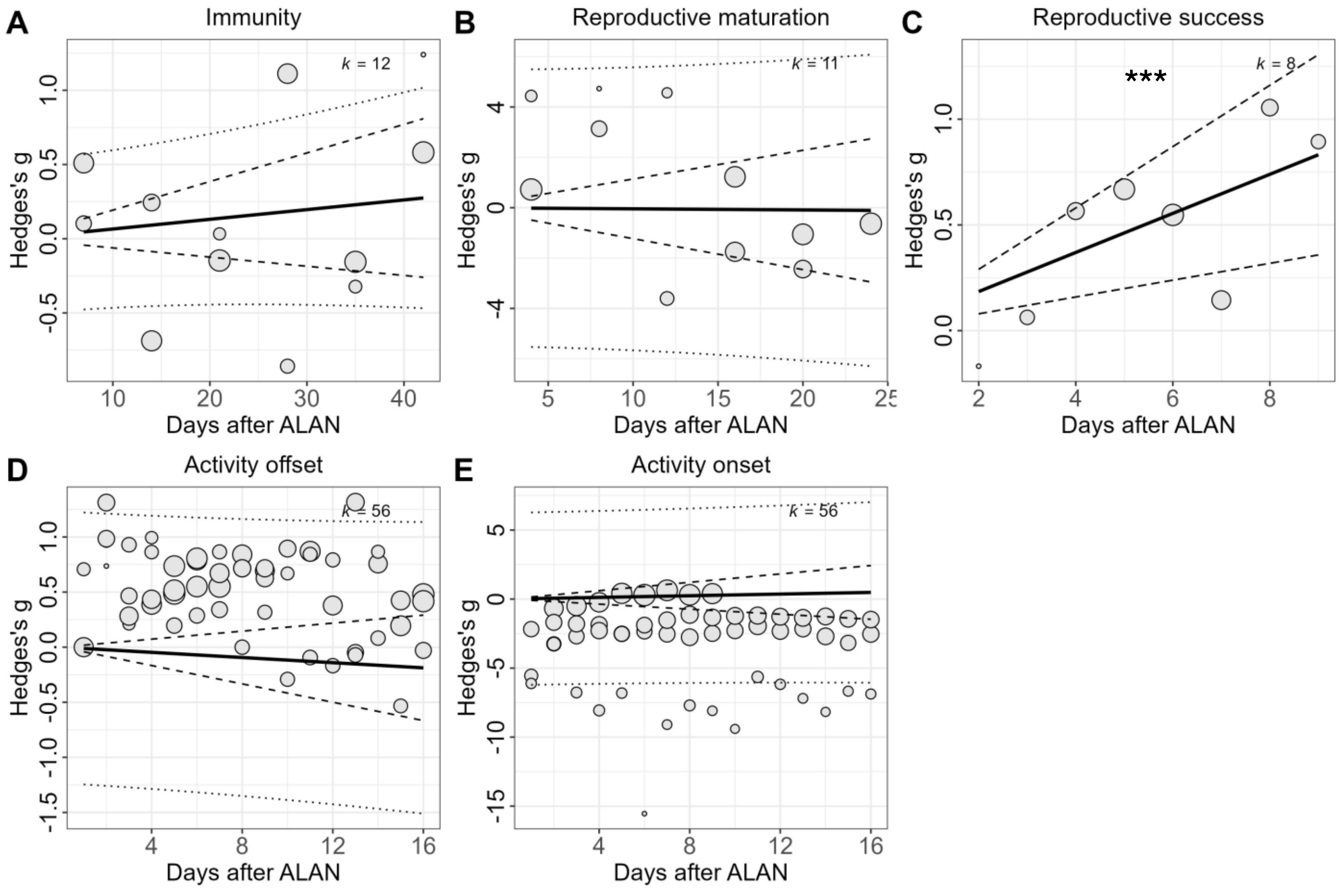


**Figure S4.** Results from the model 13 testing the impact of days after exposure to artificial light at night (ALAN) on avian functional traits: A) immunity B) reproductive maturation C) reproductive success, D) activity offset and E) activity onset. We ran a separate model for each functional trait that included repeated effect sizes measured at different days after ALAN. Longer exposure to ALAN (more days) was associated with significantly higher reproductive success, while the remaining functional traits were not affected by the time passed after ALAN exposure. The bubble plots show Hedges' effect sizes on the y-axis, and days after ALAN exposure on the x-axis. The solid line represents the model estimate, with 95% confidence intervals shown as dashed lines and 95% prediction intervals as dotted lines. The asterisks *** show the significance level $P < 0.001$ and *k* represents the number of effect sizes for each level. Full model 13 outputs are shown in Table S6.

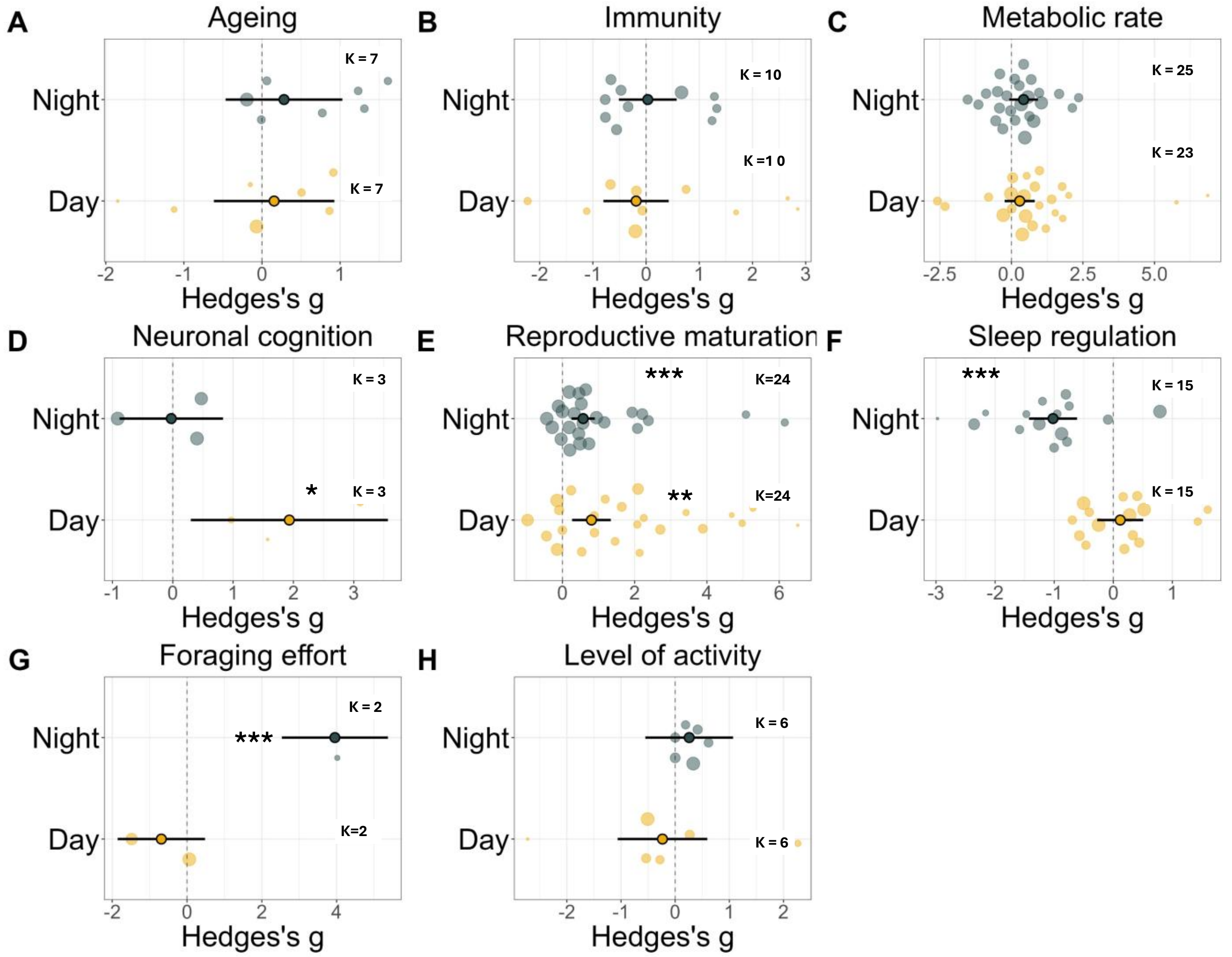


**Figure S5.** Impact of light pollution on avian functional traits for day and night time points. A) ageing B) immunity, C) metabolic rate, D)neuronal cognition, E) reproductive maturation, and F) sleep regulation, G) foraging effort and H) level of activity. The orchard plots show model estimates for Hedge's g (x-axis) along with its 95% CIs (thick whisker) and 95% prediction intervals (thin whisker). Positive and negative estimates indicate positive and negative effects of ALAN, respectively. Results from meta-analytical models including time of day (day or night) as moderator. Day (yellow) and night (grey) time points are displayed in the y-axis. The asterisks show the significance level (*P < 0.05, **P < 0.01 and ***P < 0.001) and k represents the number of effect sizes for each level. Full model 14 outputs are shown in Table S6.

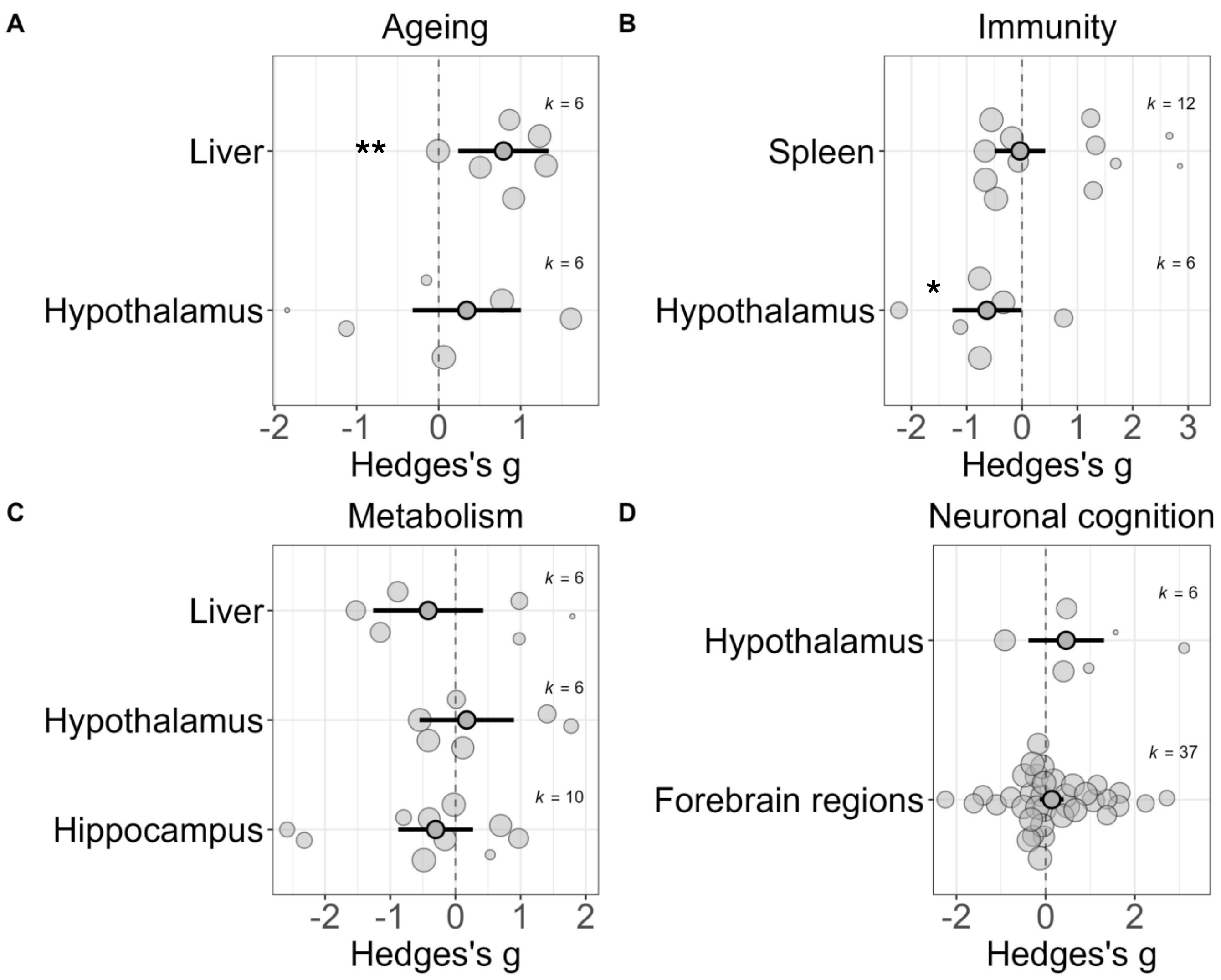


**Figure S6.** Tissue-specific effects of artificial light at night (ALAN) on avian functional traits. Each plot represents a model for a specific functional trait: A) ageing B) immunity, C) metabolic rate, and D) neuronal cognition. ALAN significantly accelerated ageing in the liver and reduced immunity in hypothalamus, while the rest of functional traits did not show any significant effects. The orchard plots show model estimates for Hedge's g (x-axis) along with its 95% CIs (thick whisker) and 95% prediction intervals (thin whisker). Positive and negative estimates indicate positive and negative effects of ALAN, respectively. Results from meta-analytical models including tissue type as moderator. The asterisks show the significance level (*P < 0.05, **P < 0.01 and ***P < 0.001) and k represents the number of effect sizes for each level. Full model 15 outputs are shown in Table S6.

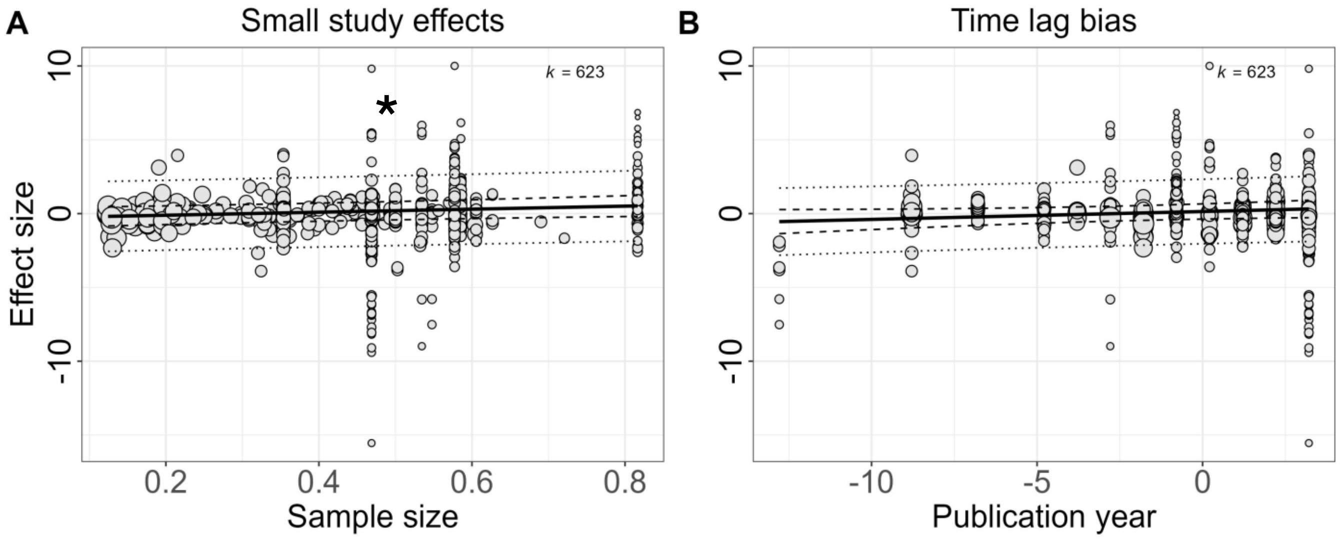


**Figure S7.** Results of Model 3 testing for publication bias on the size of the effects found in the dataset used for the overall model. A) Small study effects model showed significant effects indicating that studies with smaller sample sizes have larger treatment effects than those with larger sample sizes. The study sample size was transformed using the square root of the inverse of the sample size (x-axis) with the effect size (y-axis). B) Time lag bias model testing if more statistically significant effects are published quicker than smaller showed no significant effects. The mean-centred of the publication year was included in the model as moderator and is shown in (x-axis). In both bubble plots, the solid line represents the model estimate with the dashed line representing the 95% confidence intervals and the dotted lines representing the 95% prediction intervals. The asterisks show the significance level (*P < 0.05) and k represents the number of effect sizes for each level.
